# Filtered Point Processes Tractably Capture Rhythmic And Broadband Power Spectral Structure in Neural Electrophysiological Recordings

**DOI:** 10.1101/2024.10.01.616132

**Authors:** Patrick F. Bloniasz, Shohei Oyama, Emily P. Stephen

## Abstract

Neural electrophysiological recordings arise from interacting rhythmic (oscillatory) and broadband (aperiodic) biological subprocesses. Both rhythmic and broadband processes contribute to the neural power spectrum, which decomposes the variance of a neural recording across frequencies. Although an extensive body of literature has successfully studied rhythms in various diseases and brain states, researchers only recently have systematically studied the characteristics of broadband effects in the power spectrum. Broadband effects can generally be categorized as 1) shifts in power across all frequencies, which correlate with changes in local firing rates and 2) changes in the overall shape of the power spectrum, such as the spectral slope or power law exponent. Shape changes are evident in various conditions and brain states, influenced by factors such as excitation to inhibition balance, age, and various diseases. It is increasingly recognized that broadband and rhythmic effects can interact on a sub-second timescale. For example, broadband power is time-locked to the phase of <1 Hz rhythms in propofol induced unconsciousness. Modeling tools that explicitly deal with both rhythmic and broadband contributors to the power spectrum and that capture their interactions are essential to help improve the interpretability of power spectral effects. Here, we introduce a tractable stochastic forward modeling framework designed to capture both narrowband and broadband spectral effects when prior knowledge or theory about the primary biophysical processes involved is available. Population-level neural recordings are modeled as the sum of filtered point processes (FPPs), each representing the contribution of a different biophysical process such as action potentials or postsynaptic potentials of different types. Our approach builds on prior neuroscience FPP work by allowing multiple interacting processes and time-varying firing rates and by deriving theoretical power spectra and cross-spectra. We demonstrate several properties of the models, including that they divide the power spectrum into frequency ranges dominated by rhythmic and broadband effects, and that they can capture spectral effects across multiple timescales, including sub-second cross-frequency coupling. The framework can be used to interpret empirically observed power spectra and cross-frequency coupling effects in biophysical terms, which bridges the gap between theoretical models and experimental results.

---

Electrical field recordings in the brain – such as local field potentials (LFP), electrocorticography (ECoG), or electroencephalography (EEG) – are composed of rhythmic and broadband signal information [1, 2]. This information can be quantified by estimating the power spectrum, which decomposes the variance of a signal across frequencies [3, 4, 5]. Since field potentials themselves are driven by the superposition of many underlying subprocesses [6, 7], the neural power spectrum reflects the frequency content of these subprocesses. By separating the power by frequency, the power spectrum can reveal the effects of subprocesses that have very small magnitude at high frequencies, which would be hidden in the time domain by larger low-frequency effects. A long history of empirical work has identified spectral features that reliably correlate with brain states, originally with rhythms that show up as narrowband peaks, and more recently extending to broadband features related to the shape of power spectrum across wide frequency ranges. These rhythmic and broadband spectral features are useful tools for quantifying power spectral effects, but they provide an incomplete picture of brain states and their underlying dynamics. To better understand healthy and pathological brain dynamics measured in field potentials, it is critical to connect the features of their power spectrum to underlying biophysical processes.

Rhythmic information, sometimes referred to as periodic or oscillatory activity, has received a great deal of attention at least since Berger’s [8] introduction of the ‘alpha’ and ‘beta’ waves using EEG. Since then, rhythmic activity has been associated with a wide variety of behavioral mechanisms and brain states [e.g., 9, 10, 11, 12], and has been proposed as a mechanism for information gating within and across brain regions [e.g., 13, 14, 15]. Analyses on canonical narrowband frequency ranges (e.g., theta, 4-8 Hz; alpha, 8-12 Hz) are commonly used as biomarkers to track and compare changes in rhythms under particular brain states. Focusing on traditional power spectral bands for representing periodic signals is complicated, however, by rhythms that are transient [16, 17] or not strictly sinusoidal [18, 19]. Such complexities can lead to inconsistent reporting of spectral biomarker results across different subdisciplines in neuroscience [20, 21].

Studying rhythmic activity through power spectra is additionally complicated by the presence of broadband spectral effects, in which power across a wide range of frequencies varies together without a well-defined narrowband peak, distinct from any narrowband rhythmic effects in the signal and confounding measures of narrowband power [e.g., 22, 23, 24, 25, 26, 21, 27, 28]. Broadband effects are referred to by different names in the literature, often in a nonequivalent manner. These include aperiodic activity, power law scaling, 1/*f*-like activity, high-gamma activity, and scale-free activity [29, 30, 31, 32, 33, 34]. Here, we use the term ‘broadband’ to refer to any power spectral feature or effect without a well-defined spectral peak. At least two broadband effects clearly represented in the frequency domain have been reported as consistent biomarkers of brain state: broadband height and broadband slope.

Broadband height is typically measured by the average power in the high-gamma band (80-200 Hz), due to the rarity of narrowband rhythms in this range (c.f. sharp-wave ripples, [25, 35]). High gamma power is a reliable indicator of population firing rate [36, 37, 38, 39, 40, 41, 42, 1, 43] and dendritic activity [29, 44, 45] in the absence of rhythms. The other biomarker of the underlying network activity, spectral slope, refers to the power-law exponent required to fit a 1/*f* ^*β*^-like distribution to the measured power spectrum, i.e. the slope of the spectrum on a log-log plot. Broadband slope at moderate frequency ranges (e.g. 30-100 Hz) has been shown to change systematically in a host of conditions and brain states, including excitation to inhibition (E:I) balance [46, 47], age [48, 49, 50, 51, 52, 53], epilepsy [54, 55], autism [56], sleep [57], c.f. [58], schizophrenia [59], ADHD [60, 61, 62], and treatment-resistant depression [63]–among others (see table 1 in [64] for additional references across brain states).

**Table 1:** Figure 1 Awake State Model Parameters.

| Awake FFP Model Parameters | Value |
| --- | --- |
| Rhythm Max Power | 1500.0 *spikes/sec* |
| Rhythm Center Frequency | 13.0690611 Hz |
| Rhythm Frequency Width | 2.59384288 Hz |
| AMPA $\lambda_0$ | 9777.8253 *spikes/sec* |
| GABA $\lambda_0$ | 420.672889 *spikes/sec* |
| Scalar 1 | 977.551196 |
| Scalar 2 | 1000.0 |
| AMPA $\tau_{decay}$ | 3.3 ms |
| GABA $\tau_{decay}$ | 50.0 ms |
| AMPA $\tau_{rise}$ | 0.163508138 ms |
| GABA $\tau_{rise}$ | 0.1 ms |
| Low-pass $\tau_{LP}$ | 80.9611576 ms |

**Table 2:**
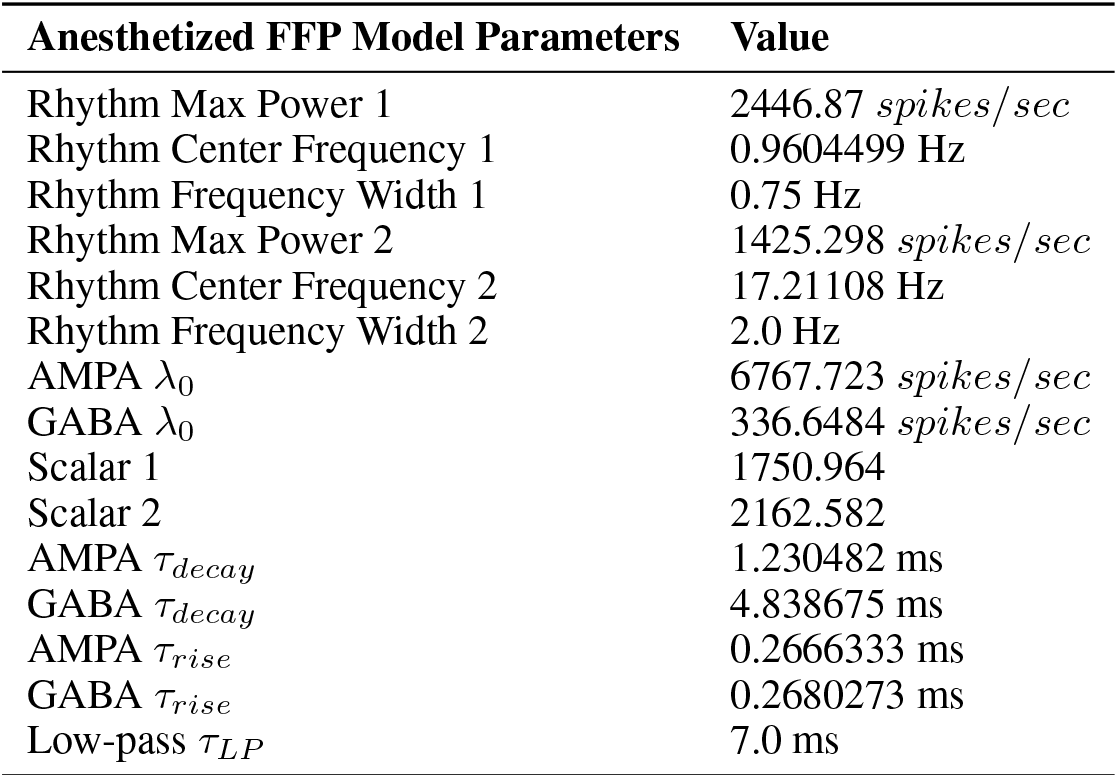
Figure 1 Anesthetized State Model Parameters.

**Table 3:** Figure 1 Infusion State Model Parameters.

| Infusion FFP Model Parameters | Value |
| --- | --- |
| AMPA $\lambda_0$ | 22767.5109 *spikes/sec* |
| GABA $\lambda_0$ | 679.678298 *spikes/sec* |
| Scalar 1 | 103.742954 |
| Scalar 2 | 243.979565 |
| AMPA $\tau_{decay}$ | 7.99997309 ms |
| GABA $\tau_{decay}$ | 9.81179409 ms |
| AMPA $\tau_{rise}$ | 0.35979477 ms |
| GABA $\tau_{rise}$ | 0.120320066 ms |
| Low-pass $\tau_{LP}$ | 6.54483556 ms |

Rhythmic and broadband features overlap in the power spectrum, can change dynamically in time, and can interact with each other. Nevertheless, there is a critical lack of theoretical models that simultaneously capture both narrowband and broadband spectral features. Traditional models of neural dynamics, whether at the single neuron, network, or population scale, typically use low-frequency time-domain dynamics and narrowband rhythmic dynamics as benchmarks for model fit, so they do not match the high frequency variance and broadband spectral shape of observed neural signals [e.g., 65]. One category of models that does address broadband effects is empirically motivated, decomposing narrowband and broadband effects using functional regression or Gaussian processes [e.g., 66, 67, 68, 69, 70]. For example, ‘specparam’ (formerly ‘FOOOF’) fits the broadband shape as log-linear with one or more knees to capture changes in the slope, and fits rhythms as bumps added to the broadband background [68]. The tool has been applied to establish spectral biomarkers across sleep, anesthesia, cognitive tasks, and clinical contexts and has undergone independent methodological study [34]. While such spectral decomposition approaches capture both narrowband and broadband effects, they are intended for descriptive analytics and they therefore do not attempt to model their biophysical generators. This limits the interpretative value of identified rhythmic and broadband components. For example, the same underlying biological processes can have both narrowband and broadband effects on the power spectrum, leading to interactions between the components that are not captured by existing parametric spectral decomposition techniques. To address this limitation, we propose a complementary approach: using forward modeling to capture how narrowband and broadband spectral effects arise from underlying biophysical generators, how they mix together in the observed power spectra, and how they interact with each other dynamically.

To capture both rhythmic and broadband effects, we will build from an existing class of models that focuses on broadband effects in the absence of rhythms. These models explain broadband effects as arising from changes in the rates of random events (modeled as point processes), which each have a characteristic waveform shape (modeled as a linear filter) [29, 32, 46]. We refer to these models as Filtered Point Processes (FPPs). The events in question can be action potentials, leading to the prediction that higher population firing rates will cause higher broadband power at particular frequencies, but they can also be other electrical events that occur with random timing. Many of the biggest contributors to neural fields are event-like in nature: in addition to action potential events, postsynaptic potentials are thought to often be the largest contributing waveform to cortical LFP recordings [e.g., 6, 71, 72], and dendritic calcium events have been shown to be detectable on the cortical surface [45, 73].

The intuition for how random event times can create broadband spectral effects is as follows. A single, perfectly sharp spike occurring once in a time series (a delta function) has power across all frequencies, i.e. it has a flat power spectrum. A series of multiple spikes at random times also has power across all frequencies, and the amount of power is proportional to the number or rate of the spikes. If we have a series of events that are not spikes but that occur at random times, for example a series of postsynaptic potentials, the spectrum will also have power at all frequencies but it will not be flat – there will be more power at lower frequencies because the smooth waveforms act like a low-pass filter. However, the height of the power spectrum at any given frequency will still be related to the rate of the events. If the rate of events increases or decreases, the whole power spectrum will shift up and down, which is a broadband effect. Furthermore, the waveform shape will determine the shape of the power spectrum, and if there are multiple event types with different waveform shapes, their superposition can lead to complex shape and slope effects in the overall power spectrum. While this perspective can capture broadband shape effects, previous work has not attempted to additionally capture rhythmic or low-frequency effects arising from dynamics in the event rates.

Here, we generalize the original filtered point process models [e.g., 29, 32, 74] to account for a wider array of rhythmic and broadband power spectral effects. We build on previous literature in at least four ways. First, we model rhythmic or time-varying firing rates. Second, we allow for the presence of multiple interacting subprocesses, for example excitatory and inhibitory populations with dynamic coupling. Third, we derive the theoretical power spectra for the models at both long and short timescales. Finally, we show how the resulting forward modeling framework can predict important qualitative effects in neural power spectra, including transition points between rhythmically-dominated vs broadband-dominated regions of the power spectrum, and cross-frequency coupling at sub-second timescales. By relating proposed biophysical subprocesses to their dynamic narrowband and broadband effects on the power spectrum, our framework will enable practitioners to test candidate models of the biophysical sources of changes in spectral shape and cross-frequency coupling.

## Theory

Filtered point process models start from the idea that the biophysical generators of neural voltage recordings are mostly characterized by discrete events, such as local action potentials, postsynaptic potentials, and dendritic spiking events [6, 7]. In other words, continuous neural signals are generated by processes that may be modeled as point processes. The event-like nature of the processes underlying the recordings is not always clear because each event is not a perfect spike, but rather has a characteristic temporal shape or waveform: if the rate of events is very high, the contribution of each event will be obscured in the time domain. Nevertheless, the event rates and the shapes of their waveforms will have predictable consequences on the power spectrum of the signal.

FPP models take the perspective of a recording electrode in space, modeling the field potential as the superposition of the voltage response generated by subprocesses representing several different event types. Figure 1 shows an example of this approach, describing how the shape of the power spectrum of surface ECoG recordings in macaques changes under propofol anesthesia (data from [75], model extended from [46]. See Supplemental Material for details). The observed power spectrum (Panel A) changes dramatically from the awake state to the unconscious state. We model the signal as a superposition of two subprocesses (Panel B), related to the excitatory synapses (red) and inhibitory synapses (blue) on the pyramidal cells underlying the electrode. Each subprocess is modeled as a random time series of postsynaptic events, filtered by the characteristic postsynaptic current shape (Panel B, insets). By formally modeling this structure, we can derive the theoretical power spectrum of the combined model (Panel C). Each subprocess contributes two components to the power spectrum: one component related to the time-varying rhythms or ensemble dynamics (solid red/blue lines) and one component related to the broadband effects of random event times (dashed red/blue lines). The theoretical power spectrum of the model (black line) is a combination of these components.

**Figure 1:**
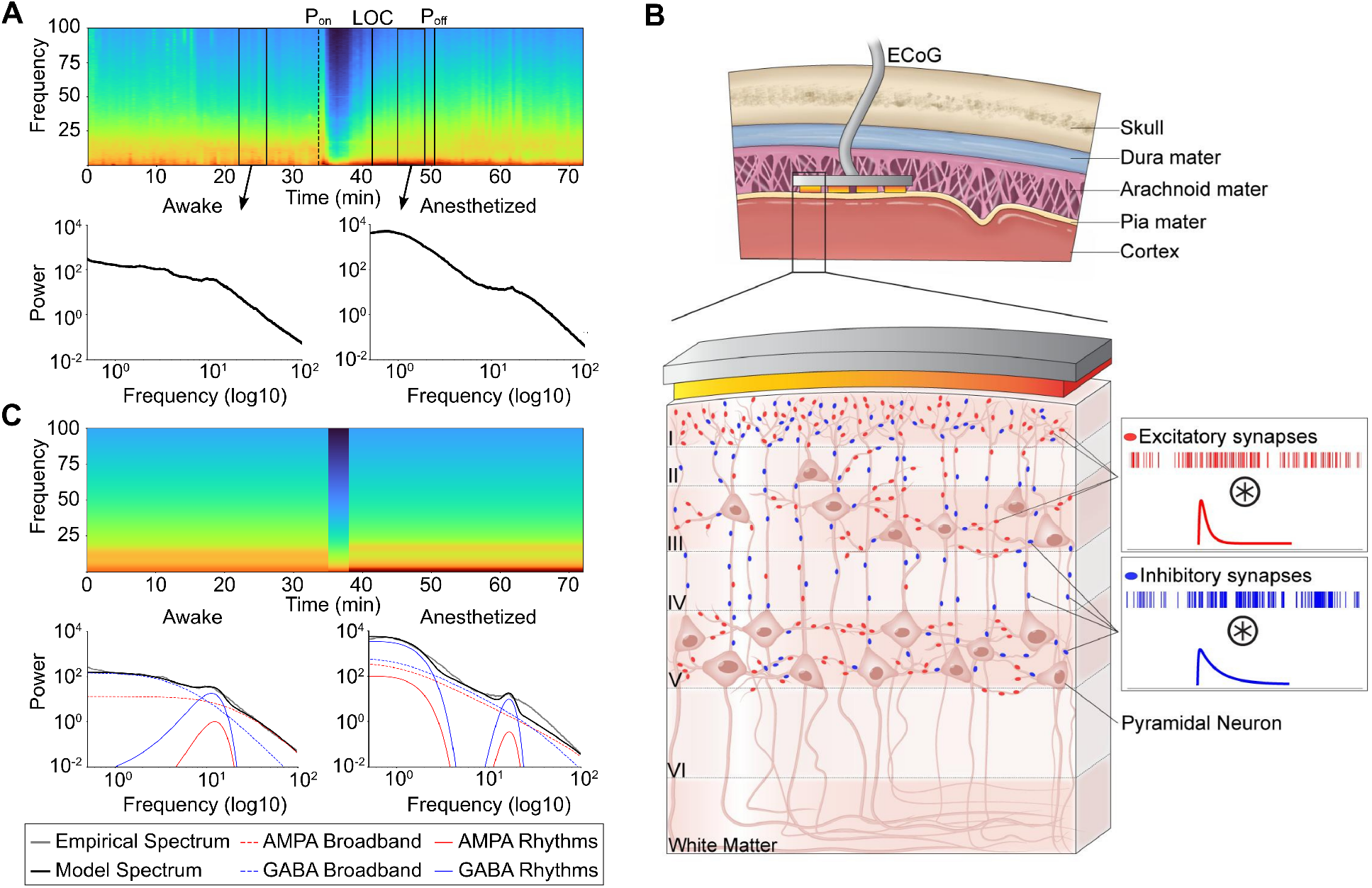
Filtered point process models describe field recordings as a superposition of event type subprocesses. Shown is a motivating use case for the filtered point process (FPP) framework: modeling the influence of excitatory and inhibitory synaptic dynamics on surface voltage recordings. (A) Spectrotemporal analysis of electrocorticography (ECoG) recordings from a monkey undergoing propofol anesthesia using data from [75]. (Above) Spectrogram of the entire session (*P*_*on*_: propofol onset; *LOC*: loss of behavioral consciousness; *P*_*off*_ : propofol offset) (Below) Mean power spectra from the Awake and Anesthetized (behaviorally unconscious) states. (B) FPPs model multiple subprocesses; here, excitatory (red) and inhibitory (blue) synaptic events on the pyramidal cells underneath the electrode. Each subprocess has a stereotyped waveform occurring at random event times (insets). (C) Example FPP models of the awake and anesthetized states shown in Panel A. The models allow a decomposition of the power spectra into rhythmic (solid lines) and broadband (dashed lines) components. See Supplemental Material for data and modeling details.

The example in Figure 1 Panel C illustrates how models of this form can be used to test the ability of proposed mechanisms to explain power spectra and support hypothesis generation. In the awake state, the LFP spectrum across all frequencies is well described by postsynaptic potentials [6], with an AMPA-GABA mediated alpha rhythm [8, 76, 77] and with inhibitory activity dominating low frequencies [7]. In the unconscious state under propofol, classic theory would include a slow wave and an alpha rhythm [76, 78, 79, 80, 77], with a steeper spectral slope at high frequencies due to increased inhibition [46]. When we include just those elements in this model, it fails to describe the power between ∼30 and 100 Hz. This could motivate us to iteratively adjust the model, for example with a beta-gamma rhythm [81] or with alternative broadband effects [82, 83, 28]. Below, we will describe previous work with related models and then present the details of our modeling framework.

## Related work: Filtered homogeneous Poisson process modeling

Filtered homogeneous Poisson point processes [FHPP; e.g., 84] are a special instance of the more general FPP framework in which the event rates do not change in time. FHPPs have emerged as an approach for modeling both temporal and spectral information in field potentials in the absence of narrowband oscillations [e.g., 29, 32, 74, 85, 46, 86, 87, 28], and they illustrate how FPPs can be used in both theoretical and applied neuroscience research. In FHPPs, each subprocess only contributes a broadband component to the total power spectrum, and the overall spectral shape reflects the relative firing rates of the subprocesses. FHPPs have been used to justify the relationship between population firing rates and the broadband height of the power spectrum [24, 1, 23, 25, 35, 85], as well as how excitatory-inhibitory balance can influence the slope of the power spectrum [46]. These theoretical studies used forward modeling and simulation to postulate the generators of observed effects, and they have motivated applied research, such as in brain-computer interfaces [36, 37, 38, 39, 88], where high gamma power (80-200 Hz) is a common proxy for population firing rates. The success of these models in both theoretical and empirical research speaks to their substantial explanatory power, despite the simplicity (and linearity) of their structure.

Despite the value of the FHPP models, they do not allow for narrowband effects like rhythms, because they assume a constant firing rate. However, trial-based experiments and many brain states have complex firing rates and oscillations that vary as a function of time. Further, prior models have not accounted for multiple interacting subprocesses, which would be necessary to model coherent rhythms. By generalizing FPPs to allow time-varying and coherent firing rates, we extend their explanatory power by capturing more features of observed power spectra.

## Model Structure

The FPP framework has a modular structure (Figure 2), building up a model based on the (1) event timing and (2) event waveforms for the subprocesses of interest, with (3) the option to add additional filters to capture effects such as low-pass filtering due to properties of the extracellular medium. A given subprocess represents the information from many different sources (e.g., synapses) from many different neurons in the vicinity of the electrode. In the next few sections, we will build up a model example using Figure 2, where Panels C, F, and I show the time and frequency representations of an example model.

**Figure 2:**
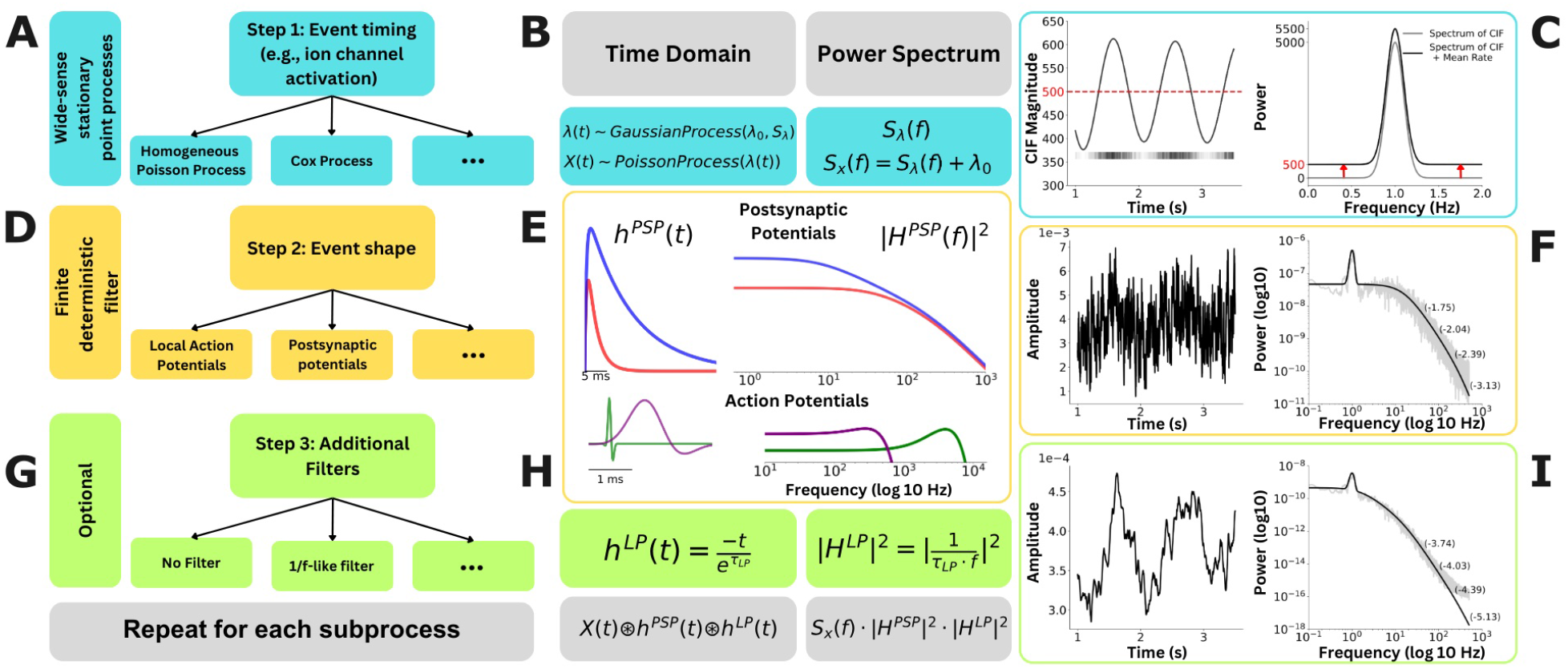
The filtered point process (FPP) framework. (Row 1: A-C) Event time series with optionally dynamic rates. (A) A wide variety of point processes can be integrated into the FPP framework provided that the point process itself is wide-sense stationary. (B) One desirable example is the Cox process, a point process with a random CIF, here modeled by a spectrally-defined Gaussian Process. (C) (Right) The theoretical spectrum of an example CIF (gray) with peak power at 1 Hz and a peak width of 0.1 Hz; the point process spectrum (black) is shifted up by 500 (the mean rate) on a linear scale. (Left) A realization of the CIF, with mean 500 *spikes/sec*. The raster plot below shows the point process event times. **(Row 2: D-F) Event waveform filters** (D) Any linear, finite, and deterministic filter can be used in the FPP. (E) Several filter templates have been derived. Top: two postsynaptic potentials *h*^*PSP*^ (t), AMPA (red) and GABA (blue), defined in the time domain, and *H*^*PSP*^ (f), capturing the effects of the filters on the power spectrum. Bottom: two action potentials (AP), regular spiking (green) and fast spiking (purple). (F) The effect of a GABA filter on the point process example from Panel C: (Left) the effect on the time-domain realization; and (Right) the effect on the theoretical (black) and empirical (gray) power spectra. Slopes of the theoretical power spectrum at four points are annotated in parentheses. **(Row 3:G-I) Optional additional filters** (G) Additional filters can be used to capture different shape effects driven by the system being modeled. (H) (Left) A generic low-pass filter specified in the time domain, and (Right) its effect on the power spectrum in the frequency domain. Below in gray is the theoretical solution for the full model in the time and frequency domain. (I) Results of the final model for the local field potential (Left) in the time domain and (Right) the theoretical (black) and empirical (grey) power spectra of the process, with four example slopes annotated in parentheses.

## Event Timing

The basis of the model are a set of event types in the neural tissue underlying the electrode, such as postsynaptic potentials, which are considered to be important contributors to the rhythms and/or broadband power in the spectrum. The event *times* are modeled by point processes (Figure 2, teal), capturing the structure of the randomness in the event times [89]. In general, point processes are parameterized by a Conditional Intensity Function (CIF) λ(*t*), which captures the instantaneous rate of events; note that any point process can be fully specified by its CIF, which is given as the joint probability density of event times [89]. For example, homogeneous Poisson processes assume that the CIF is constant in time 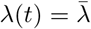. By allowing the CIF to vary randomly in time, we can account for dynamic variation in the mean-field or ensemble dynamics of the event type. This is achieved by a Cox process, a point process where the CIF is itself a stochastic process [84, 89, 90, 91], making the overall point process “doubly stochastic” (i.e. the CIF is random and the event times are random conditioned on the CIF).

Figure 2 Panels B and C show an example of a Cox process that can capture a slow oscillation at 1 Hz (details in Methods). The time-varying CIF is modeled by a Gaussian process with mean λ_0_ and power spectrum *S*_*λ*_(*f*), where the power spectrum contains a narrowband peak at 1 Hz (Panel C, gray line). The point process is then modeled as an inhomogeneous (i.e. time-varying) Poisson process where the rate λ(*t*) is the realization of the Gaussian process:

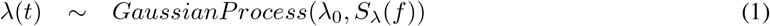

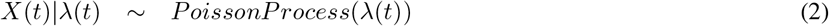

where *X*(*t*) is a time series representing the random event times of the Poisson process. Here we assume that the mean rate λ_0_ is large enough that the fluctuations in the CIF λ(*t*) are highly unlikely to be negative. This assumption is often justified by the large number of event generators (e.g. synapses, cells) in the vicinity of a recording electrode. In cases where the assumption is unlikely to hold, a link function can be used to guarantee non-negative event rates, such as a log link function [5].

We can also represent *X*(*t*) by its counting process, *N*_*k*_, which represents the number of spikes in discrete bins, where bin *k* is defined as the interval from time *k*Δ*t* and time (*k* + 1)Δ*t*. The number of events in any time bin is Poisson distributed, and the rate is approximated by the CIF at time *k*Δ*t*:

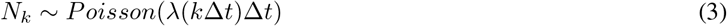

for small Δ*t*. In neuroscience recordings, it is common to choose Δ*t* = 1*ms*, and represent *N*_*k*_ as a 0-1 vector in discrete time. The framework is flexible with regard to discrete and continuous representations of time.

By definition, the power spectrum of a stochastic process is the Fourier transform of the autocovariance function (dependence across time lags) of the process, so it fully describes the second moment of the process in the frequency domain [3, 4, 5]. Spectral analysis assumes that the process is weak-sense stationary, meaning that the first moment (mean) and second moment (autocovariance) do not vary in time within the time window that is being analyzed. Here, because the time-varying λ(*t*) is governed by a stationary Gaussian process, the overall point process is also stationary. Hence, the process (and by extension the full FPP model) has a well-defined power spectrum.

We would like to understand the structure of the power spectrum of the point process *X*(*t*), since it will form the basis of the power spectrum of the full filtered point process. A technical detail here is that point processes use a generalized notion of a power spectrum, the Bartlett spectrum. For our purposes, it is sufficient to know that the Bartlett spectrum behaves like a standard spectrum under filtering [90]. The Bartlett spectrum of *X*(*t*) contains two terms, one related to the power spectrum of the CIF and one related to the mean rate [92, 90]:

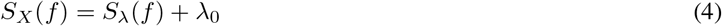

The mean rate of events (λ_0_) controls the height of the spectrum at all frequencies. Temporal dependence in the CIF (rhythms, autocovariance) will lead to variations in the spectrum across frequencies (*S*_*λ*_(*f*)).

Figure 2 Panel C shows how this works for the slow wave Cox process example: on the left, a realization of the process has a CIF (solid black line) that oscillates at approximately 1 cycle per second and an event timeseries (black raster plot) where events are more likely to occur when the CIF is high than when it is low. On the right, the theoretical power spectrum of the Cox process (black line) is the sum of the power spectrum of the Gaussian process (gray line) and a flat component equal to the mean rate (here 500 *spikes/sec*). We will refer to this flat component as the “broadband” component of the power spectrum.

## Event Shape

Each point process is filtered with an event shape or waveform, which corresponds to the extracellular waveform of the biological process of the corresponding event type (Figure 2, yellow). These can range from action potentials, which may account for as much as 20*%* of the electrical field [72], to postsynaptic potentials, which are thought to make up the bulk of local field potentials [93, 6, 7]. Waveforms can be flexibly defined, and their effect on the theoretical power spectrum is analytically tractable whenever the Fourier transform is well-defined. We refer to the time domain representation of events as “waveform,” “filter,” or “event shape,” whereas a “kernel” is the magnitude squared of the Fourier transform of the waveform. The filter information can be combined with point process information in the time and frequency domains through a convolution in the time domain or multiplication in the frequency domain, retaining the tractability of the solution.

Figure 2 Panel E shows four example waveforms: two postsynaptic potential types (AMPA and GABA, in red and blue), and two action potential types (regular spiking and fast spiking, in green and purple; see Methods for details).

For postsynaptic potentials, we use the difference of two exponentials as a generic functional form, which can be tuned by its rise and decay time constants to capture the dynamics of different channel types [46, 71, 86, 94]. Let *h*^*PSP*^ (*t*) be the shape of the postsynaptic potential in the time domain:

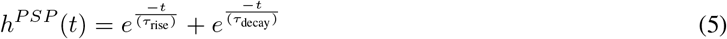

where *τ*_rise_ and *τ*_decay_ are the rise and decay time constants of the postsynaptic currents, which will vary depending on the type of synapses (i.e., AMPA and GABA synapses).

For the action potential waveforms, we use a real Gabor function (c.f. [86]), which can be tuned to create fast and slow action potentials:

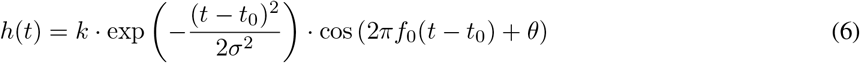

where *k* is the amplitude of the action potential, *f*_0_ is the center frequency of the cosine function, and *θ* is the phase offset.

For any event type, the filter *h*(*t*) is applied to the point process *X*(*t*) in the time domain via a convolution:

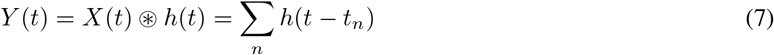

where *t*_*n*_ are the times of the events in *X*(*t*).

This convolution in the time domain will correspond to a multiplication in the frequency domain (i.e., a Fourier pair). If we let ℋ(*f*) be the Fourier transform of *h*(*t*), then the effect of the filter will be to multiply the power spectrum of *X, S*_*X*_ (*f*), by the magnitude squared of ℋ [3]:

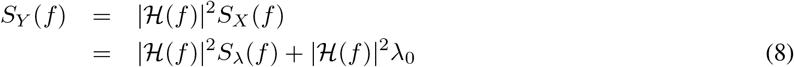

Notice that the overall power spectrum decomposes into two terms, inherited from the two terms in Equation 4: the first term relates to the dynamics of the event rates, and the second term relates only to the mean rate of the events. While both terms are shaped by the frequency-dependence of the filter, the first term will typically be narrowband, while the second term will typically be broadband. Equations 7 and 8 hold for any wide-sense stationary point process and finite, deterministic filter selection.

The waveforms in Panel E have well-defined Fourier transforms, which can be used to compute the kernel |ℋ (*f*)|^2^ and applied to Equation 8 to derive the theoretical power spectrum of the FPP. For the postsynaptic potentials (Equation 5), the Fourier transform is:

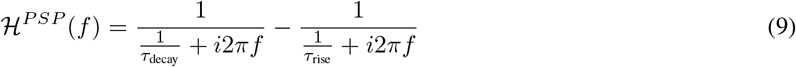

And for the action potentials (Equation 6), the Fourier transform is:

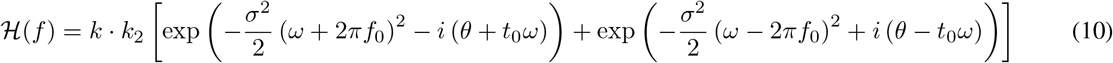

where *ω* = 2*πf* is the angular frequency and *k*_2_ is a distinct scaling constant (see Supplement 4 for full equation).

The framework can also be used in conjunction with empirically fit waveforms, such as action potential shapes that are known to vary across cell types [95, 96, 97, 98, 99, 100]; in this case, a discrete Fourier transform can be used to compute the kernel of the empirical waveform.

Figure 2 Panel F shows the effects of applying a GABA waveform to the time-domain realization and theoretical power spectrum of the Cox process from Panel C. In the time domain realization (left), the waveform transforms the time series from a sequence of events into a noisy continuous-time signal, where the 1 Hz rhythmic structure in the event rates is visible in the trend. In the power spectrum (right), the spectral peak from the CIF is still present, and the broadband component has been shaped by the function |ℋ^*PSP*^ (*f*)|^2^, which is approximately flat for low frequencies and curves down with a noticeable negative slope above 10 Hz. In fact, for the postsynaptic potential shape shown in Figure 2 Panel E and used in the model for Panel F, the slope of the filter will converge to *β* =−4 in log-log space as frequency tends to infinity (see Supplemental Materials for derivation). However, as shown in Figure 2 Panel F, this limiting slope does not apply to these models until frequencies above those typically examined in neuroscience studies.

## Additional Filters

While the properties of event timing and waveform shape play a large role in the shape of the power spectrum on an extracellular electrode, there may be other effects that need to be modeled using additional filters – either for theoretical reasons or to achieve a better match to empirically observed data.

For example, additional filters could capture the transformation of transmembrane currents to extracellular potentials, based on the biophysical properties of the cell membrane and extracellular medium [101, 93, 71, 102]. The underlying properties of extracellular media and their effect on the frequency scaling of field potentials continue to be an active area of research. As such, we include this unknown in our modeling framework by including the possibility of additional filters that can be designed to represent the desired biophysical effects.

Additional filters can also be helpful to match empirically observed data. An empirical effect that has been of particular interest because of its ubiquity across spatial scales and recording modalities is the power-law scaling property of field potential power spectra. “Power-law scaling” or “1/*f* ^*β*^ frequency scaling” refers to the empirical observation that the power spectrum often forms a line in some frequency ranges when plotted on log-log axes, i.e. the power in log space is inversely proportional to the frequency f increased to an exponent *β* [103, 32, 64]. The exponent *β* is referred to as the “spectral exponent” or “spectral slope”, and empirical spectral exponents of neural power spectra vary between about -2.5 and -4 across a variety of healthy and diseased brain states [64]. As shown in Figure 2 Panel F, an FPP model with only a postsynaptic potential filter has slopes between zero and -3.1 for frequencies below 1000 Hz. Additional theoretically justified low-pass filters may be helpful to capture steeper slopes.

For example, a simple low-pass filter representing the effects of ionic diffusion can be given as:

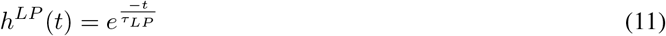

where *τ*_*LP*_ is the time constants of the filter. The Fourier transform of the filter is:

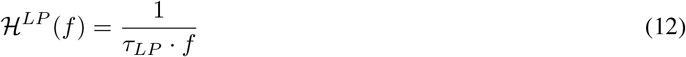

When applied to the model with postsynaptic potentials (Equations 7), the filter operates as a convolution in the time domain:

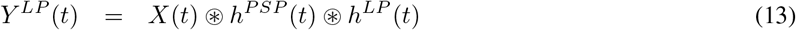

and the power spectrum (Equation 8) is shaped by the filter kernel:

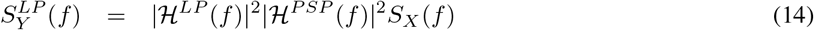

Figure 2 Panel I shows the impact of this low-pass filter on the signal from Panel F. In the time-domain realization (left), the filter smooths out much of the noise in the signal, and the 1 Hz oscillation is more apparent. In the power spectrum (right), the filter has tilted the entire power spectrum, making the slopes more negative, although the original 1 Hz spectral peak is still visible. Here, the the low-pass filter itself has a spectral slope of *β* =−2 and, when combined with the spectrum in Panel I, the whole model will have a limiting slope of *β* =−6 (see Supplemental Materials for derivation). Ultimately, the final power spectrum captures both periodic information, through the dynamics of the CIF, and broadband information, through the stochastic point process.

## FPPs can model multiple subprocesses simultaneously

When multiple subprocesses are modeled, for example both AMPA and GABA postsynaptic potentials, the field recording is considered to be the superposition or sum of the filtered subprocesses:

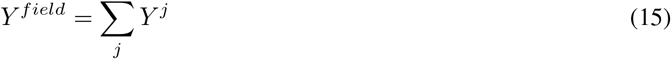

where *Y* ^*j*^ are the filtered subprocesses, for example described by Equation 13.

In this case, it is important to model the dynamic dependence of the subprocesses, because coherence between subprocesses will affect power spectrum of the field recording. For example, if we have two point processes relating to AMPA and GABA postsynaptic potentials, we can model their CIFs using a bivariate Gaussian process, and the point processes as a bivariate Poisson process conditioned on the rates from the Gaussian process:

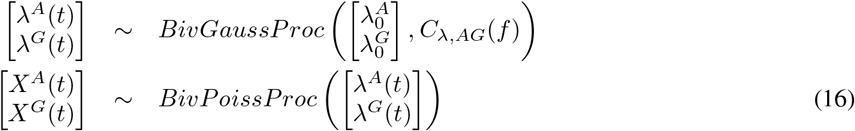

where *A* refers to the AMPA subprocess, *G* refers to the GABA subprocess, and λ_0_’s are the overall mean rates. For each frequency *f, C*_*λ,AG*_(*f*) is a 2 *×* 2 cross-spectral matrix of the CIFs, which can be written as:

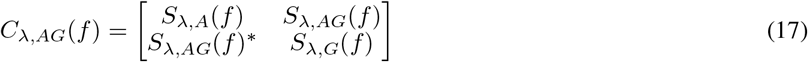

where *S*_*λ,A*_(*f*) and *S*_*λ,G*_(*f*) are the power spectra of the AMPA and GABA CIFs at frequency *f*, respectively, and *S*_*λ,AG*_(*f*) is the scalar cross-spectrum (the Fourier transform of the cross-covariance function) between the two CIFs at frequency f. This captures the coherence structure between the two CIFs, because (complex) coherence is defined as the normalized cross-spectrum:

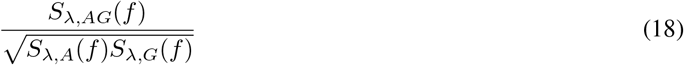

We assume that the only dependence between the processes is via the CIFs, i.e. that the actual event times are conditionally independent after accounting for the CIFs. If the two CIFs are independent, then *S*_*λ,AG*_(*f*) = 0 for all frequencies.

Understanding the effect of dependence between subprocesses on the FPP model amounts to describing how each model choice affects the cross-spectrum of the resulting process. First, the cross-spectrum of the bivariate point process is given by [89]:

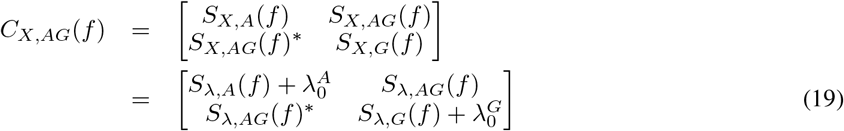

We can see from these equations that the mean rates 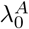 and 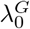 only affect the diagonal terms (the power spectra), not the dependence between the two processes (the cross-spectrum).

Applying filters to the model will shape both the spectra and the cross-spectrum. For example, if we filter *X*^*A*^(*t*) by *h*^*A*^(*t*), the AMPA waveform, and *X*^*G*^(*t*) by *h*^*G*^(*t*), the GABA waveform:

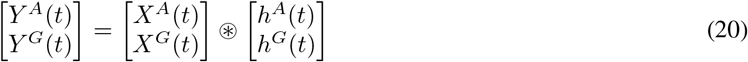

then the cross-spectral matrix will be (suppressing the dependence on *f* for compactness)[3]:

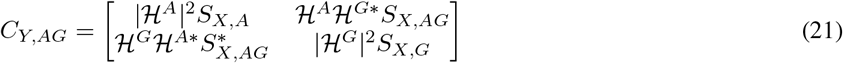

where, as before, ℋ^*A*^(*f*) and ℋ^*G*^(*f*) are the Fourier transforms of *h*^*A*^(*t*) and *h*^*G*^(*t*), respectively. We can see that the spectra on the diagonals are transformed as expected from Equation 8, and the cross-spectrum is shaped by the product of the kernels or the Fourier transforms of the AMPA and GABA waveforms. This step can be repeated for any additional desired filters.

Finally, the power spectrum of the final field recording *Y* ^*field*^, as defined in Equation 15 will be:

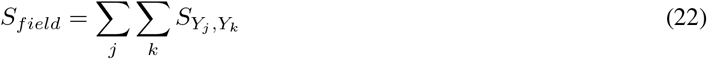

or in the AMPA and GABA example:

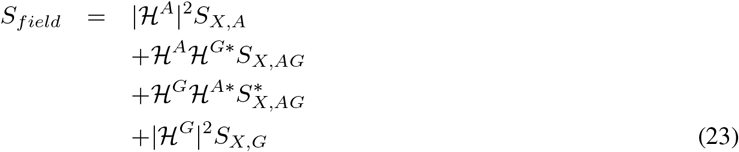

Additional filters that are applied equally to all components represented in *S*_*field*_, such as the low pass filter |ℋ^*LP*^ (*f*)|^2^ introduced in Equation 14, can simply be represented as:

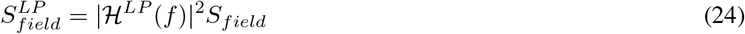

This would then be distributed as

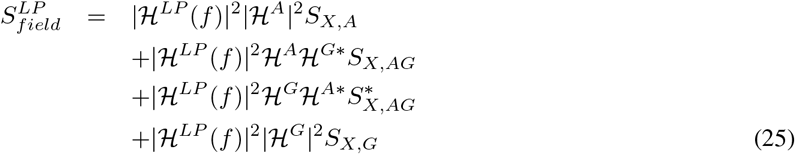

We now present two special cases of this bivariate FPP structure with AMPA and GABA synapses: independent subprocesses and processes that share a CIF up to a linear transform.

First, if the processes (including their CIFs) are independent, then the off-diagonal elements of the cross-spectral matrix (Equation 17) are zero for all frequencies, so the final field spectrum will simply be the sum of the spectra of the two processes:

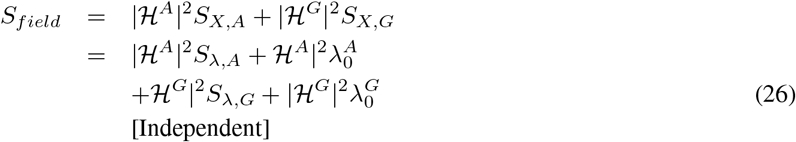

If the processes share a CIF up to a linear transform, i.e. if:

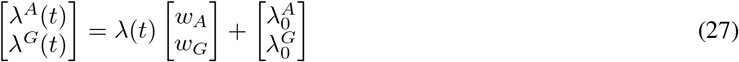

where λ(*t*) is a univariate Gaussian process with spectrum *S*_*λ*_(*f*) and 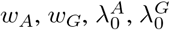 are known parameters, then the power spectrum of the CIF will have a term for the spectrum of the CIF, shaped by the two kernels, and two terms related to the mean rates of the processes:

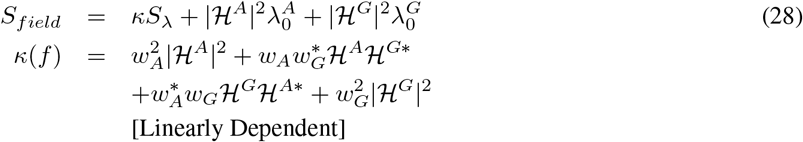

Hence, as in the case with only one subprocess (Equation 8), the power spectrum for a model with multiple subprocesses decomposes into terms related to the spectra of the CIFs (typically narrowband) and terms related to the mean rates (typically broadband).

## Results

### Result 1: FPP power spectra are dominated by either ensemble dynamics or broadband at different frequencies

Power spectral effects in neural voltage recordings tend to show frequency ranges that are dominated by either rhythmic or broadband effects — this is the basis for defining canonical rhythms (e.g. slow <1 Hz) and fixed bands for broadband power (e.g. high gamma 80-200 Hz). Empirically, rhythms tend to dominate at lower frequencies and broadband effects tend to dominate at higher frequencies. Rhythmic or not, the lower frequencies can be dominated by coherent afferent projections to the recorded area [e.g., 104, 105, 106]. Signals traveling longer distances tend to exhibit slower temporal dynamics, which lend themselves well to synchronization (e.g., [107]). In contrast, higher frequency components are correlated with local firing behavior, as described above [108, 105, 32, 42, 109]. FPP models capture and explain this property of neural power spectra: frequencies will either be dominated by “ensemble dynamics”, i.e. the power spectrum of the rate process (CIF), or “broadband”, i.e. the flat power spectrum arising from random event times. This is a direct result of the power spectrum of a point process given in Equation 4.

In Figure 3, we demonstrate this effect using a FPP model with a single GABA subprocess for simplicity. The mean rate λ_0_ and the spectrum of the CIF *S*_*λ*_(*f*) are shown in Figure 3 Panel A, green and blue lines, respectively. The mean rate component is “broadband” because it has constant power across all frequencies. If the modeled point process is a homogeneous Poisson process, this broadband component is the only spectral information present. When a FPP is modeled using a Cox process, its power spectrum is additionally affected by the power spectrum of the CIF. We refer to this component as the “ensemble dynamics” in a neuroscience context because it describes the temporal structure of the rate process for the event type, e.g. the dynamic rate averaged across all GABA synapses in the ensemble. In Figure 3, the CIF is modeled as an AR(2) process that has a spectral peak.

**Figure 3:**
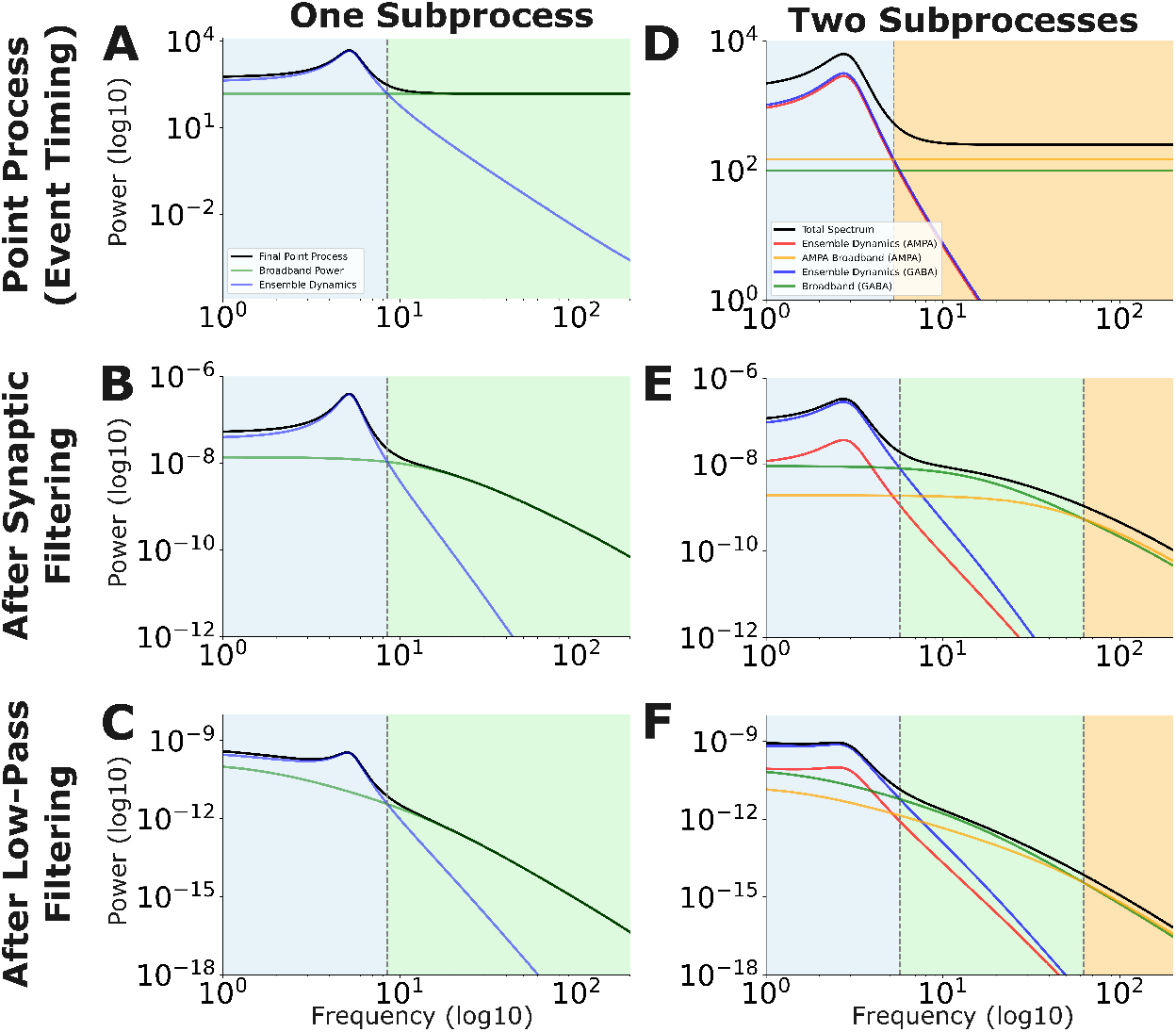
Filtered point process (FPP) models exhibit transition points in their spectra. *Column 1*: A GABA subprocess is modeled as a Cox process with an AR(2) CIF (blue line) and a mean rate (green). The overall theoretical spectrum for each modeling step is given in black. (A) The mean rate component and the spectrum of the conditional intensity function (CIF) produces a transition point ∼7.3 Hz. This produces two regions of the power spectrum, CIF-dominated (blue shading) and broadband-dominated (green shading) region. Applying the GABA filter (B) and a 1/f-like low-pass filter (C) changes spectral shape but does not change the transition point. *Column 2*: Two independent AMPA and GABA subprocesses are modeled, each with AR(2) CIFs. The overall theoretical spectrum for each modeling step is given in black. (D) The CIFs of the AMPA and GABA process (unfiltered) are shown in orange and blue, respectively, and their broadband components are in red and green. Shading reflects the dominant component, as before, and the transition point is ∼5.27 Hz. (E) Filtering the subprocesses by different filters (the AMPA and GABA event shapes) changes the transition points. The final spectrum now has two transition points: GABA ensemble dynamics dominate below ∼5.7 Hz, GABA broadband dominates from 5.7-62.65 Hz, and AMPA broadband dominates above 62.65 Hz. E) Spectra after applying a 1/f filter: the transition points do not change.

For any single subprocess modeled as a Cox FPP, both the ensemble dynamics and broadband information have the potential to contribute at any frequency. However, FPP models predict that certain components of the model will dominate at certain frequencies, leading to overall power spectral shape effects. Figure 3 Panel A shows the power spectrum of a point process with an AR(2) CIF. The overall power (black line) at frequencies shaded in blue primarily reflects information contributed by the ensemble dynamics (blue line) and the power at frequencies shaded in green primarily reflects information contributed by the broadband (green line). Around 7.3 Hz, there is a transition point: ensemble dynamics power dominates below 7.3 Hz and the broadband power dominates above 7.3 Hz. This pattern of different components dominating at different frequencies has been observed before in other filter-based models [110, 28]. Given the parameters of the ensemble dynamics, there will be a transition point as long as the mean rate line crosses the CIF line.

Importantly, the location of the transition point does not change when the process is filtered: Figure 3 Panel B shows the power spectrum after applying the GABA postsynaptic potential shape to each event via filtering, and Panel C applies an additional 1/f-like filter. It can be seen that the transition point is identical after filtering, i.e. any given frequency is dominated by either ensemble dynamics or broadband power, and the choice is determined at the level of the point process, regardless of any subsequent filtering.

This has several practical implications for interpreting empirical neural power spectra. First, according to this model, rhythmic effects and broadband effects are not additive in log space, as some spectral decomposition methods assume [68, 111]. Rather, the effects are additive in linear space [112, 66, 70], causing the largest component to dominate when viewed on a log scale. Second, broadband shape effects such as the spectral slope or power law exponent will be effectively invisible at frequencies that are dominated by rhythms or other ensemble dynamics. To estimate the spectral slope it is therefore necessary to restrict analysis to frequencies that are dominated by broadband effects. Note that in some cases, the power law exponent of the ensemble dynamics may be of interest [113, 49, 114, 115, 116] in which case it will be important to restrict analysis to frequency ranges dominated by those effects.

### Result 2: Multiple FPP subprocesses can produce complex spectral shape effects explained by simple model components

Parsimonious descriptions of field potentials will typically highlight at least two subproccesses as the primary drivers of a neural recording. FPPs can accommodate this by modeling multiple subprocesses with either independent or dependent temporal structure. Here, we focus on two independent subprocesses, process 1 (ultimately filtered by an AMPA waveform) and process 2 (ultimately filtered by a GABA waveform), and show that the components of the subprocess power spectra themselves have transition points that can be useful in explaining complex power spectral shape effects.

Figure 3 Panel D shows two independent point processes modeled with nearly identical ensemble dynamics modeled as AR(2) processes (process 1 in orange and process 2 in blue). Additionally, process 1 and process 2 have two different broadband components, shown in red and green respectively. As with the previous section, each subprocess has one transition point between the ensemble dynamics and broadband. Overall, lower frequencies are dominated by the process 2 ensemble dynamics and the higher frequencies are dominated by process 1 broadband, so the highlighted transition point is between two different subprocesses.

When introducing the AMPA (process 1) and GABA (process 2) filters in Panel E, more complex relationships arise. First, as expected, the transition point within each subprocess is preserved (green/blue crossing and red/orange crossing). However, there is a change in the overall dominant component from applying different filters to each subprocess. While the GABA ensemble dynamics still dominate at low frequencies (blue shading in Figure 3), in this model the addition of the postsynaptic filters allows the GABA and AMPA broadband components to cross and create a region where GABA broadband dominates (shaded green) and a region where AMPA broadband dominates (shaded orange). Depending on the kernel structure, this can produce relatively complex shape effects beyond slope changes.

Finally, even with multiple subprocesses, applying an identical 1/f-like filter across all subprocesses does not change the transition points between processes (Figure 3 Panel F).

This result shows that, under the FPP modeling regime, there is far more information represented in a neural power spectrum than merely narrowband oscllations or changes in 1/f-like slope in an otherwise uniform aperiodic spectral component. Rather, different frequency regions of the spectrum could reflect different subprocesses that have their own CIF and broadband components. In fact, it makes the prediction that the power spectrum could represent the emergence of different subprocesses dominating on different timescales, such as in the case of up- and down-states, which we explore in Result 4.

### Result 3: Subprocess transition points are determined by the relative magnitude of ensemble dynamics and broadband

In Result 1, it was shown that, given fixed ensemble dynamics and broadband parameters, transition points and dominant components do not change when filtering for a single subprocess. In Result 2, it was shown that additional transition points can emerge at the level of modeling subprocesses. Here, we show that changing the relative magnitude between model components at the level of a single subprocess shifts where transition points can occur–both within a subprocess and across interacting subprocesses.

As a case study, we show a single GABA subprocess with an AR(2) CIF in Figure 4 Panel A; it has already been filtered by both the GABA and 1/f-like filter and has a transition point established around 6.56 Hz. There are two ways that the transition point can shift: the broadband component can increase or decrease in power relative to ensemble dynamics or vice versa. In Panel B, the ensemble dynamics are kept constant and the broadband component is increased from 50 to 400 *spikes/sec*. This results in a shift of the transition point from around 6.56 Hz to 4.39 Hz, substantially increasing the impact of the broadband on the overall power spectral shape. Likewise, in Panel C, keeping the AR(2) coefficient the same and decreasing the AR(2) Gaussian (state) noise from 0.315 to 0.1 which induces a vertical and linear transformation across all frequencies, we see a shift from around 6.56 Hz to around 4.23 Hz. As such, relatively small changes in the relative magnitude of model components can have a large impact on the shape and interpretability of the power spectrum for given frequencies. Similarly, while shifting either component can produce comparable changes in the actual frequencies at which a given component is dominating the power spectrum, the final power spectra are different in their overall interpretation.

**Figure 4:**
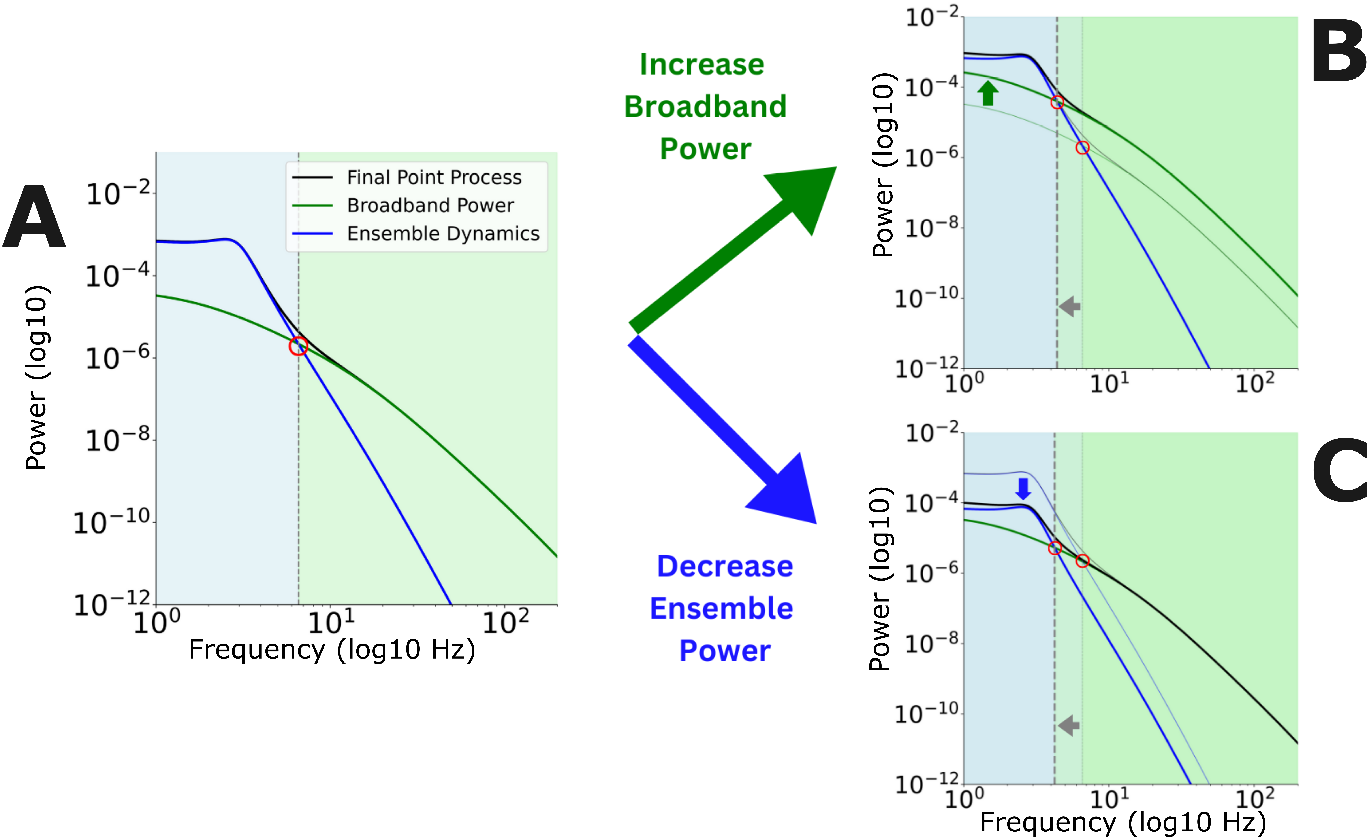
Transition points between model components are determined by the relative magnitude between model components. A) Shown are the spectral components of a Cox process with an AR(2) conditional intensity function (CIF) and a mean rate. The dominating component in the blue region is the spectrum of the CIF (blue line) and the dominating component in the green region is the mean rate (green line). The transition point takes place at ∼6.56 Hz. B) Keeping the spectrum of the CIF constant, an increase in the mean rate of the process moves the transition point to lower frequencies (4.39 Hz). Both transition points are circled in red and their relative movement of the transition points is highlighted by the gray array. The relative movement of the mean rate is highlighted by the green arrow. C) Keeping the mean rate of the process constant, a decrease in the power of the CIF spectrum (blue arrow) results in a shift of the transition point to lower frequencies (4.23 Hz). Movements in transition points suggest that different frequency regions of the power spectrum, even in the absence of rhythms, should be interpreted in a time-dependent manner with respect to the underlying dynamics.

As a result, FPP models suggest that rhythmic information across brain states or tasks most likely is not contained in a fixed, relevant frequency band. Rather, the frequencies at which rhythmic or broadband information dominates can change drastically depending on the relative magnitudes of the ensemble dynamics versus broadband components. As such, tracking narrowband oscillation changes (e.g., traditional power analyses) or changes in frequency domain functional connectivity (e.g., Granger-Geweke causality analysis) across different global brain states or task structures could be ambiguous without using forward modeling to generate expectations for how frequencies of interest could change. This is consistent with previous arguments presented in [19, 68, 2]. Furthermore, it suggests that smoothly shifting oscillations (e.g., drug infusion-induced changes) are not necessarily driven by changes in the mechanism of an oscillation itself but could be due to a stable oscillation that shifts in presentation as the broadband effects change. Forward models that can accurately account for both rhythmic and broadband information can be used to rule out such possibilities given knowledge of the underlying mechanisms.

### Result 4: FPP models capture both overall spectral structure and fine-timescale dynamics

As described in the Theory section, FPP models are stationary, meaning that their statistical properties do not vary in time. This is one of the key features allowing for the full tractability of the modeling approach, which yields overall (i.e. long-timescale) theoretical power spectra. Nevertheless, for any given realization of the process, the instantaneous rate of events can be dynamic, i.e. the CIF λ(*t*) can vary in time. If we condition on λ(*t*), the resulting point process describing event times, *X*(*t*|λ(*t*)), is an inhomogeneous (i.e. non-stationary) Poisson process. We can use this property to study changes in the expected shape of the power spectrum on short timescales.

To study short-timescale dynamics, we will need to assume that λ(*t*) is smooth or slowly varying, so that the rate of events is approximately constant in short time windows. This property is called semi-stationarity, and it is a common assumption when studying neural data using spectrograms [117, 118, 119]. By semi-stationarity, we can approximate the inhomogeneous Poisson process in a short time window by a homogeneous (i.e. constant-rate) Poisson process. We can then use the homogeneous Poisson process as the event process in our FPP model, deriving the theoretical power spectrum for that time window.

As a case study of this property, Figure 5 shows a bivariate Cox FPP model, derived in Equation 28, with AMPA and GABA subprocess components. The theoretical power spectrum (Panel A) shows the long timescale spectral shape, where the GABA subprocess dominates the power at frequencies below around 16 Hz and the AMPA subprocess dominates for higher frequencies. This is consistent with previous simulation work looking at AMPA and GABA processes interacting during propofol-induced unconsciousness [46], except here we’ve accounted for an additional narrow band slow wave rhythm that was not included in the prior work. We model the slow wave using a single Gaussian process with a spectral peak at 1 Hz: the AMPA and GABA CIFs multiply this process by 1.5 and -0.8, respectively.

**Figure 5:**
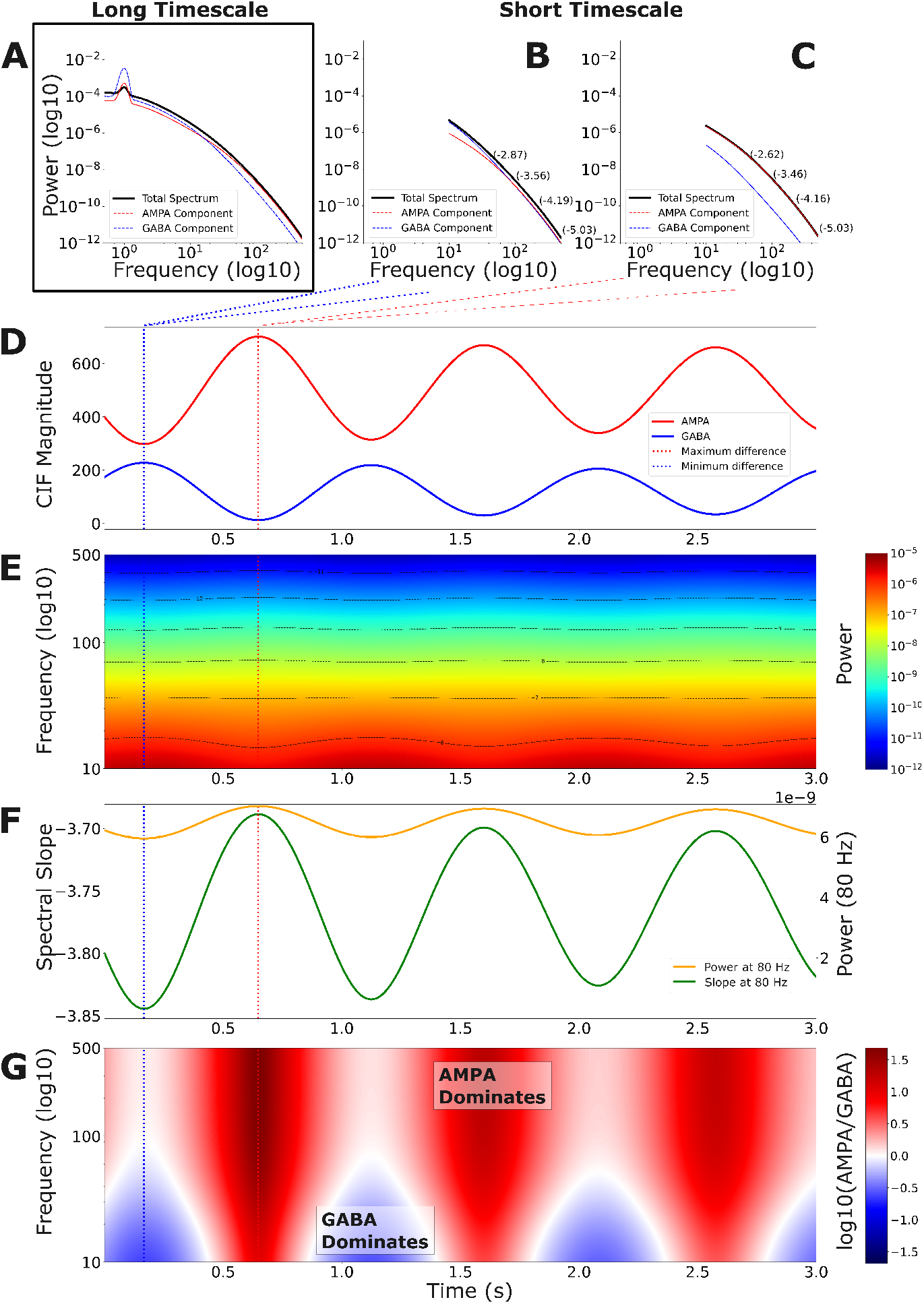
Filtered point processes (FPPs) capture distinct information at different timescales. (A) The theoretical power spectrum of a field potential modeled as a bivariate Cox process with AMPA and GABA subprocess components and a 1 Hz slow wave oscillation over a long timescale. The CIFs of each subprocess are identical up to a column vector that scales each CIF. (D) A single realization of the scaled CIFs for each subprocess for AMPA and GABA, respectively, and are then offset by the mean rate of each process. Assuming the realized conditional intensity function is semi-stationary on a short timescale, we can approximate higher frequencies of its power spectrum theoretically as a homogeneous Poisson process. (B) and (C) Two extremes where the AMPA and GABA CIFs are minimally and maximally distant in magnitude, respectively. This makes the prediction that the short-time power spectra will have periods where the overall spectral shape is dominated by primarily AMPA dynamics, whereas others will have lower frequencies dominated by GABA and higher frequencies are dominated by AMPA. Additionally, the spectral slope values at 40, 90, 200, 450 Hz for the total spectrum are shown, indicating that these models have not yet converged to the overall model limiting slope (see Supplemental Materials for derivation). (E) These semi-stationary dynamics at a fine timescale can be visualized across total time in the continuously varying theoretical spectrogram. (F) The spectral slope and power at 80Hz as a function of time. (G) The times and frequencies at which a given subprocess component is dominating the spectral shape can be highlighted by showing log ratio of AMPA and GABA components from the short-time spectrogram.

We can study short-timescale properties in this model by generating a realization of the CIFs (Panel D), treating the process as semi-stationary, and investigating the properties of the time-varying theoretical power spectrum (Panels B, C, E, F, and G). In Panel D, it can be seen that the CIFs for AMPA and GABA oscillate at about 1 Hz, out of phase: the AMPA subprocess has an up-state when the GABA subprocess has a down-state and vice versa.

The semi-stationarity assumption allows us to study theoretical short-time power spectra at different time points (Panels B and C). At the peak of the AMPA process and the trough of the GABA process (Panel C), the overall spectral shape is dominated by AMPA power. At the trough of the AMPA process and the peak of the GABA process (Panel B), in contrast, frequencies below about 93 Hz are dominated by GABA power and higher frequencies are dominated by AMPA power. Hence the transition point between AMPA-dominated and GABA-dominated frequencies varies dynamically according to the phase of the rhythm, and even disappears when AMPA rates are high.

Note that the short timescale theoretical power spectra in Panels B and C are limited to frequencies above 10 Hz, while the long-timescale spectrum in Panel A has frequencies below 1 Hz. While we were able to compute these spectra at a single time point using the instantaneous model parameters, they are not truly instantaneous power spectra. Rather, they relate to time windows long enough for the CIFs to be “approximately constant”. Here, the CIFs vary at about 1 Hz, so we assume that windows of about 100 ms have approximately constant rates: since the lowest resolvable frequency in a 100 ms window is 10 Hz, we truncate the power spectra at 10 Hz.

Also shown in Panels B and C are the spectral slope values at several selected frequencies. Importantly, the spectral slope is not the same across frequencies – for very high frequencies it converges to -6 (see Theory and supplemental derivations), but the slopes in the frequency range observed here have not converged to this limiting value. In other words, spectral slopes at frequencies typically studied in neuroscience research may have less to do with a theoretical limiting “power law”, and more to do with the filter shapes of the component subprocesses and their relative magnitudes.

From these short-timescale power spectra, a theoretical spectrogram (Panel E) can be generated to build an intuition for how the resulting process would look as a function of time, changing dynamically as the balance of excitation to inhibition changes. By looking at the contours, we can see that the 1 Hz oscillation is affecting low and high frequencies differently: for low frequencies, power is higher when the GABA power is higher, while for high frequencies, power is higher when AMPA power is higher. In the middle, around 40 Hz, is a frequency were power is not affected much by the oscillation. In traditional terms, the spectral slope and high frequency broadband power are both changing dynamically during the rhythm (Panel F shows the spectral slope and power at 80 Hz).

The source of these effects is the changing relative contributions of the AMPA and GABA components: to facilitate interpretation, we show the time varying log ratio of the AMPA and GABA components (Panel G). This highlights the times and frequencies when the power spectrum is dominated by the AMPA component (red) or the GABA component (blue). We can also see the transition point between AMPA-dominated and GABA-dominated frequencies as a white band, varying dynamically from about 93 Hz at the peak of the GABA process to disappearing at the peak of the AMPA process. Hence we see from this example how the broadband components and the rhythm can *interact*: here, both the height and the shape of the overall power spectrum vary as a function of the rhythmic phase. For more complicated models, the changes in power spectral shape at short timescales could be even more complex.

This analysis of timescales makes a strong prediction, which is that cross-frequency coupling between rhythmic phase and broadband power should be ubiquitous in neural electrophysiological recordings. Under the model, every rhythmic component is accompanied by a broadband component for the same subprocess, and on short timescales the height of the broadband component will change dynamically with the phase of the rhythm. Of course, the broadband components may be obscured by larger components, for example when the GABA broadband component is obscured by the AMPA broadband component at the peak of the AMPA rhythm in Figure 5. In some situations, strong non-rhythmic broadband components may completely hide any cross-frequency coupling in weaker subprocesses. Nevertheless, this result predicts a very close relationship between rhythms and broadband power (and shape) that could be instrumental in interpreting cross-frequency coupling in biophysical terms.

Practically, this type of timescales analysis can be used to validate theoretical models of neural dynamics in comparison to data. For example, FPP models of rhythms can specify which synapse types participate in the rhythms at which phases; then long- and short-timescale model predictions can be compared to empirical spectra and spectrograms, using the entire spectral shape to assess model performance. Rhythmic-broadband interactions vary at sub-second or low-second timescales in multiple brain states [e.g., 120, 121, 122, 123]. This phenomenon has been called “spectral-switching” [e.g., 121, 122, 123]; FPPs could serve as a strong theoretical framework for studying the biophysical generators of these effects.

## Discussion

In this paper, we have presented a tractable probabilistic forward modeling framework that can explain rhythmic and broadband information in neural power spectra using filtered point processes (FPPs). Building on prior work modeling broadband effects in the power spectrum using random event times filtered by characteristic event shapes, we introduced dynamic event rates to account for rhythmic and time-varying ensemble firing rates, and we introduced multiple interacting subprocesses to account for different synapse types or populations in the modeled tissue. We showed that the theoretical power spectrum of the modeled field recordings remains tractable, and that it decomposes into narrowband and broadband components related to each subprocess (Equations 8 (one subprocess) and 23-28 (multiple subprocesses)). The resulting framework can be used to evaluate whether proposed biophysical mechanisms can explain observed spectral effects, including both rhythmic and broadband effects and their dynamic relationships.

### Insights and takeaways

Analysis of representative FPP models demonstrated that analyzing power spectral shape effects as a whole provides more accurate insights into the underlying biophysical dynamics of a recording than focusing on specific features or frequency ranges. In an FPP model, broadband and ensemble contributions to the power spectrum of a subprocess are modeled explicitly. This reveals transition points in the subprocess dominating the power spectrum. Thus, the modeling question shifts from needing knowledge of all relevant system information to modeling what has the largest impact on the power spectrum at a given frequency.

For a single subprocess, the transition point between the broadband component and the ensemble component is not affected by the event shape or subsequent filtering (Result 1), but it is affected by the relative power of the ensemble dynamics and the broadband (Result 3). For multiple subprocesses, event shapes become important: faster shapes (e.g. excitatory postsynaptic potentials or action potentials) will allow processes to dominate higher frequencies, while slower shapes (e.g. inhibitory postsynaptic potentials) will constrain processes to lower frequencies (Result 2), subject to the relative firing rates of the processes. Once the event shapes are fixed, additional filters do not change transition points. The transition points within and between subprocesses can also change dynamically in time (Result 4), and treating the process as semi-stationary can facilitate analysis of the spectral properties at short (e.g. sub-second) timescales. In other words, theoretical models of rhythms and the entrainment of population firing rates could be tested with FPPs by making predictions of how power should be distributed across frequencies on both short and long timescales and comparing the predictions to empirical data.

These observations have important implications for researchers interpreting neural power spectra. First, the rhythmic and broadband components (and their interactions) add together to create the total power spectrum: this challenges popular spectral decomposition methods that decompose the power spectrum in log space, which effectively assume that the components have multiplicative effects on the power spectrum [68, 111].

Secondly, this additive relationship means that when viewed in log space, rhythmic dynamics and broadband will dominate the spectrum at different frequencies, and the spectral slope of the broadband component will be effectively invisible when analyzing slope in frequency regions dominated by rhythms. Thus, spectral slope analyses of the broadband component should be restricted to frequency ranges where there is not a known rhythm dominating the power spectrum. Similarly, there has been interest in the field about the spectral slope of low-frequency ensemble dynamics [113, 49, 114, 115, 116]: these analyses should also be restricted to frequencies where researchers do not expect other rhythms or broadband to be dominating.

Thirdly, our results caution against the use of static frequency ranges to assess the rhythmic power, broadband height, or spectral slope. Fixing predetermined oscillatory ranges (e.g., 8 - 12 Hz, 12 - 20 Hz) or broadband ranges (e.g., 80 - 200 Hz) for analysis can be misleading, since the relevant frequency ranges are likely to shift. Although this prediction would suggest that certain well-established power spectral analyses might need to be used with caution (which is consistent with previous work, e.g., [19, 20, 124, 125, 126, 127, 128]), analyzing the power spectrum as a whole provides a potentially useful perspective as a marker of changing brain states.

Finally, the FPP framework predicts that cross-frequency coupling between low-frequency rhythms and broadband spectral power and shape should be ubiquitous in neural electrophysiological recordings. If we believe that the primary contributors to the recordings are event-like (postsynaptic potentials, action potentials, etc.) and that rhythms reflect the ensemble dynamics of these events, then each rhythmic component of the power spectrum should be accompanied by a phase-locked broadband component at short timescales. While the cross-frequency coupling may be obscured by stronger dominant broadband components in some brain states, this close relationship between rhythmic and broadband power should be taken into account when interpreting cross-frequency coupling.

Altogether, our results predict that the overall shape of the power spectrum provides much more information about the underlying biophysical process than merely the rhythms, broadband height (e.g. high-gamma power) and power spectral slope. Rather, if researchers have prior knowledge about the dominant generators of a given LFP, ECoG, or EEG signal (e.g., given postsynaptic potentials, dendritic calcium spikes, local action potentials of a given cell type), which is already important for interpreting field potentials in a specific brain state [7], the entire power spectrum is interpretable beyond narrowband rhythms and a uniform broadband component [120, 121, 122, 123].

### Relationship to prior work

Neuroscience has a rich history of theoretical forward modeling, using mathematical models at the molecular, neuronal, circuit, population, and systems levels, to propose underlying mechanisms of observed dynamics and to make predictions to guide future experimental studies [129, 130, 131, 132]. In the context of modeling field potentials, conductance-based models have been used to model micro-scale extracellular potentials all the way up to scalp recordings [133, 134, 135, 136, 137]. In principle conductance-based models could account for broadband spectral effects [e.g., 138, 139] and could capture rhythmic and broadband effects simultaneously. However, as mentioned, these models often do not match the broadband spectral properties of observed neural signals [e.g., 65], because they emphasize low-frequency and narrowband rhythmic time series as model fit criteria. Modeling meso-scale broadband spectral effects using such fine-grained models is additionally difficult because of the dependence of broadband effects on the detailed spatial correlation structure across neurons, where empirical data are lacking.

There is also value in being able to generate theoretical models at higher spatial scales without the very fine level of granularity required for compartment-based neuronal modeling. Currently, two main approaches help to solve this issue: 1) single compartment neurons with conductances and 2) mean-field or neural mass models. Single compartment neurons, while often computationally tractable, often do not scale accurately to capture the empirical structure of ECoG or EEG recordings [140]. Mean-field models often focus on slow time-domain dynamics and narrowband rhythms over a mass of neural tissue [141, 142]; by design, they capture the dynamics of the population firing rate. As a result, mean-field models typically do not capture broadband power spectral features or effects because they mathematically assume the impact of autocovariance and cross-correlation on neural dynamics is negligible [82, 142]. With that in mind, mean-field models could be used in conjunction with FPP models, where they could be used in the place of the time-varying CIFs.

Filtered *homogeneous* point process models were introduced to address the limitations of previous rhythmic models and to capture broadband effects at higher spatial scales. We expect that by capturing both rhythmic and broadband components, and including multiple interacting subprocesses, FPP models will offer significantly more flexibility. The bivariate model of AMPA and GABA subprocesses has been of interest in many investigations over recent years in modeling ECoG and EEG [e.g., 46, 86, 28] and is now amenable to investigation with FPPs (Result 5). However, FPPs can also capture more general dependency between subprocesses (Equation 23).

The use case of FPP models that we described here takes the perspective of a recording electrode and models the multiple subprocesses that affect the recorded signal. As such, we have omitted literature using shot noise as input to model neurons or neural populations, which has focused primarily on fractal point processes as input to Fitzhugh-Nagumo and Hodgkin-Huxley models of peripheral nerves and neurons using renewal processes (reviewed in [143] e.g., [144, 145]. We also omit the body of literature reviewed and developed in [146], where they introduce “fractal shot noise processes,” which is a special instance of our more general framework.

Finally, we note the theoretical power spectrum for a Cox process has been well characterized in the measure-theoretic probability literature for some time [e.g., 92, 90, 91] and the theoretical power spectrum for a homogeneous Poisson process has been well characterized for use in neuroscience before [e.g., 29, 74]; however, because many recent theoretical analyses have simulated FPPs in the time domain before estimating the power spectrum nonparametrically for model analysis [46, 86], we directly provide the theoretical power spectra for a given model structure. Additionally, we provide a software package called `filtered_point_process` with simple tutorials to demonstrate how to use the framework (github repository: https://github.com/Stephen-Lab-BU/filtered-point-process).

### Limitations and Future Directions

The FPP models assume that the subprocess events sum linearly in their contribution to the recording electrode, and that the waveform shapes do not vary as a function of time or location. This is a strong assumption and clear examples exist where it does not hold. For example, postsynaptic potential shapes vary based on the time-varying intracellular potential of the electrode [130] and on the relative location of the synapse from the recording electrode [147, 148, 149]. Because our subprocesses are intended to capture many different synapses on many different neurons in the vicinity of the electrode, this could lead to significant errors in the calculation of the theoretical power spectrum. We described in the Theory section how valuable FHPP models have been to the scientific literature despite sharing this limitation. Nevertheless, in future work we plan to use nonlinear compartment-based models of cortex to evaluate the size of the effect that these approximations have on the power spectrum in specific modeled brain states. Then, we will investigate ways to alter the models to improve their performance.

For example, elegant theoretical work in integrate and fire model neurons has recently derived theoretical power spectra for the output of model neurons with nonlinear responses [150], and our framework could benefit from incorporating elements of this approach into our models. We expect that in certain brain states, it will also be important to account for variation in waveform shape across time and space. A state space model could capture time-dependence by allowing the waveform shape to change depending on the level of the CIF of the process. For the dependence on space, we could model the waveform shape as stochastic [151], as in generalized shot noise models, which could be useful for capturing variability in power spectra across a grid of electrodes (e.g., Utah arrays) driven by the positioning of a dipole source relative to the electrode position. Alternatively, it is possible to estimate effective linear waveforms across space, either theoretically using conductance-based compartment models [152, 147], or empirically as in recent work on unitary LFPs, which estimate the impact of a single axon on the local LFP [149, 100, 153]. Incorporating these empirical waveforms into the FPP framework could yield spectral estimates that are more accurate than the theoretical filters used here.

Our models do not account for refractory periods or self-history, which can have a significant effect on the power spectrum even when summed across neurons or synapses [154]. The FPP models can account for such effects by using renewal point processes or Cox processes with self-history dependence as the event time process, because the theoretical power spectra and cross-spectra of these processes are tractable [155, 156]. Interestingly, refractory periods typically have the effect of reducing low-frequency power relative to higher frequencies, regardless of the presence of low-frequency rhythms [155, 156, 5, 157], and these effects persist in the sum across neurons or synapses [154]. To our knowledge, this effect is never observed in empirical neural power spectra of field recordings, which tend to have the highest power at low frequencies. In other words, FPP models with a refractory period would likely have lower power at very low frequencies than high frequencies, which does not match empirical observations. FPP models could be built and adapted to study this paradox.

Our models are stationary, meaning that the overall properties of the models do not depend on time. While the semi-stationarity assumption can be used to study fine-timescale dynamics in the power spectrum by conditioning on the dynamic CIFs, even when the overall model is stationary, there will be some scientific questions where the stationary assumption is not appropriate. For example, different sleep stages are characterized by categorically different rhythms, and it would not make sense to use a single FPP model for all stages. In cases like these, a switching state-space approach could be used in which a discrete hidden state (e.g. the current sleep stage) controls which subprocesses and rhythms are in effect. Switching state space models have been used to great effect in modeling non-stationary power spectral effects in neuroscience [67, 158, 69].

Finally, the FPP framework is currently a probabilistic *forward* model and does not provide methods for parameter estimation, i.e. we do not yet describe how the model parameters can be estimated statistically from data. The general model framework does not guarantee that any particular model is identifiable, so there may not be enough information in a dataset to fit any particular model. In the future, we will describe principles for designing identifiable FPP models for specific brain states or neural dynamics that are simple enough to be fit to empirical data, and we will present methods for fitting such models. We believe FPPs will be a useful statistical model for empirically driven spectral decomposition, and our future work will develop the estimators, goodness-of-fit metrics, and ground-truth validation required for a successful statistical framework.

By bridging the gap between micro-scale biophysics and macro-scale narrowband and broadband biomarkers, our modeling framework enables theoretical modeling of the rapidly growing collection of known narrowband and broadband changes in healthy and diseased brain states. These theoretical models, and the software packages implementing them, can be used to improve the interpretability of neural power spectra for scientific and clinical progress.

## Materials and Methods

The FPP framework described above allows for time-varying rates by modeling the CIF λ(*t*) of each process as a Gaussian process with a known power spectrum *S*_*λ*_(*f*) (Equation 1 for a single process and Equation 16 for multiple subprocesses). We use two different types of Gaussian processes: (1) spectrally-defined Gaussian processes, where the process is defined by its mean and power spectrum (first and second moments), and (2) autoregressive (AR) processes, where the process is defined based on its dependence on previous time lags. We present each below, and then describe how each is used in the example models used in the Results.

### Spectrally-defined Gaussian Process CIFs

For spectrally-defined Gaussian processes, we choose the power spectrum of the desired CIF *S*_*λ*_(*f*). While the spectrum of a Gaussian process can be any well behaved shape, we chose to represent narrowband rhythms as Gaussian shapes or bell curves. In order to allow for multiple rhythms, the spectrum of the process is defined as having an arbitrary number of Gaussian shapes:

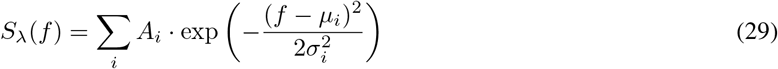

where *f* frequency; the *i*-th Gaussian peak is defined by a mean *µ*_*i*_ (center frequency), standard deviation *σ*_*i*_ (frequency width), and amplitude *A*_*i*_ (peak height).

### Autoregressive Process CIFs

Another type of Gaussian process that can be useful for modeling the CIF is a discrete time autoregressive process, which models the current time point as a function of *p* previous time points (referred to as an AR(*p*) process) [4]:

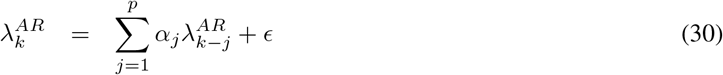

where the index *k* in 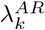 refers to the time bin, as in λ(*k*Δ*t*), and ϵ is white Gaussian noise with variance *σ*^2^. We assume that the model coefficients *α*_*j*_ are chosen so that the process is stable. The power spectrum of the process is:

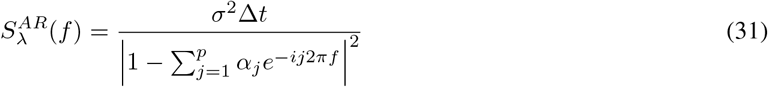

[4]. With this power spectrum and a mean rate λ_0_, we can use Equations 1-2 to define a Cox process with an Autoregressive CIF. In the case *p* = 1 the power spectrum only has a peak at 0 Hz and proceeds to decrease monotonically, which means it does not have rhythms. This case represents a discrete-time random walk, and it can be useful for modeling ensemble dynamics that vary smoothly in time without oscillating.

We generalize this result to account for multivariate AR processes. Consider the case where the CIFs obey a multivariate AR process of order 1, i.e.:

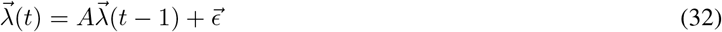

where *A* is a transition matrix and 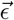 is multivariate zero-mean Gaussian noise with covariance Σ. If we assume that the process is stable, then the cross-spectrum of the process is [3]:

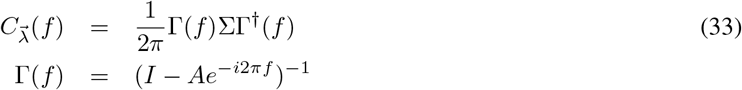

where † denotes the conjugate transpose. Note the similarity between this theoretical cross-spectrum and the spectrum of a univariate AR process given in Equation 31, where the variance *σ*^2^ has been replaced by the covariance Σ and the square in the denominator has been replaced by pre- and post-multiplying by Γ. This cross-spectrum can be used in Equations 19-23 to compute the field spectrum.

### Models used in the Results

#### Figure 2

In Figure 2, we model the CIF using a spectrally-defined Gaussian process with mean, λ_0_, of 500 *spikes/sec* and power spectrum *S*_*λ*_(*f*), where we construct the power spectrum to have a rhythm (which we modeled as a Gaussian-shaped bump using Equation 29) with width 0.1 Hz centered at 1 Hz and a height of 5000 *spikes/sec*. Defining the process using its frequency domain second moment (its power spectrum) is convenient in this context, because we focus mainly on the power spectral properties of the model. The equations for the model are specified as the following for each step: for Figure 2 Panel C, Equation 3 (left) and Equation 4 (right), for Panel F Equation 7 (left) and Equation 8 (right) where *τ*_*rise*_ = 0.4 ms and *τ*_*decay*_ = 10 ms to represent a GABA PSP, and for Panel I Equation 13 (left) and Equation 8 (right) where *τ*_*LP*_ = 10 ms.

For interpretability, the y-axis scaling in Panel C is linear in units of *spikes/sec*. This is conventional for point process power spectra [5, 157] and aids in identifying mean rate as the height of the flat component (λ_0_ in Equation 4). The y-axis scaling for Panels F and I uses the convention for continuous stochastic processes, *power*/*Hz* (e.g. *µV* ^2^/*Hz*), which differs from the point process scaling by a factor of (Δ*t*)^2^ [156, 157]. We use these conventions throughout.

The CIF described above was simulated to give a realization of Equation 3 through the approximate frequency domain method outlined in [3]. A realization of the corresponding point process was created via the standard “thinning” procedure for inhomogeneous Poisson processes [159]; this is highly efficient and is justified provided an intensity is continuous and bounded for any finite interval.

Shaded in gray in Panel F and Panel I is a multitaper estimate of 60 seconds of simulated data (a snippet is shown in the left of the figure) with a frequency resolution of 0.133 Hz. This shows that direct estimation from the simulated data correctly recovers the intended theoretical power spectrum. All parameters are summarized in table 4.

**Table 4:** Figure 2 Model Parameters.

| Model Parameters | Value |
| --- | --- |
| Rhythm Max Power | 5000 *spikes/sec* |
| Rhythm Center Frequency | 1 Hz |
| Rhythm Frequency Width | 0.1 Hz |
| GABA $\lambda_0$ | 500 *spikes/sec* |
| GABA $\tau_{decay}$ | 10 ms |
| GABA $\tau_{rise}$ | 0.4 ms |
| Low-pass $\tau_{LP}$ | 10 ms |

### Figure 3

In Figure 3 Panels A-C (Result 1), a single subprocess is modeled as a Cox FPP with a spectrum given by Equation 4. The CIF is represented as an AR(2) process, given by Equation 31. Panel A is the unfiltered point process spectrum, Panel B is the unfiltered point process spectrum shaped by the magnitude squared of a PSP filter given in Equation 9 with *τ*_*decay*_ = 10 ms and *τ*_*rise*_ = 0.4 ms, which represents a GABA kernel. Panel C multiplies the filtered spectrum in panel B by the squared magnitude of the lowpass filter given in Equation 12, where *τ*_*LP*_ = 10 ms. All parameters are summarized in table 5.

**Table 5:** Figure 3 One Process Model Parameters.

| Model Parameters | Value |
| --- | --- |
| GABA $\lambda_0$ | 150 *spikes/sec* |
| $\alpha_1$ | 1.9897557323136878 |
| $\alpha_2$ | -0.9901 |
| $\sigma^2$ | 0.7 |
| GABA $\tau_{decay}$ | 10 ms |
| GABA $\tau_{rise}$ | 0.4 ms |
| Low-pass $\tau_{LP}$ | 10 ms |

Similarly, in Figure 3 Panels D-F (Result 2) two subprocesses are modeled: process 1 and process 2. Each process spectrum is unfiltered in Panel D. In panel E, process 1 is filtered with an AMPA waveform and process 2 is filtered with a GABA waveform; additional low-pass filter was applied to each subprocess (Panel F), as was done for the single subprocess. For the GABA process, the unfiltered spectrum is given by an AR(2) process and the AMPA process is modeled as an AR(2) process, both with *fs* = 1000 Hz. All parameters are summarized in table 6. Note that the *σ*^2^ parameter is marginally offset between each process to avoid perfect overlap to demonstrate the effect of interest; the *σ*^2^ also induces a vertical shift. Panel E and F are identical to the filtering done in Panel B and C, except with the addition of the AMPA kernel given in Equation 9 with *τ*_*decay*_ = 4 ms and *τ*_*rise*_ = 0.4 ms.

**Table 6:** Figure 3 Two Process Model Parameters.

| Model Parameters | Value |
| --- | --- |
| GABA $\lambda_0$ | 100 *spikes/sec* |
| AMPA $\lambda_0$ | 150 *spikes/sec* |
| $\alpha_1$ | 1.9897557323136878 |
| $\alpha_2$ | -0.9901 |
| GABA $\sigma^2$ | 0.315 |
| AMPA $\sigma^2$ | 0.3 |
| GABA $\tau_{decay}$ | 10 ms |
| GABA $\tau_{rise}$ | 0.4 ms |
| AMPA $\tau_{decay}$ | 10 ms |
| AMPA $\tau_{rise}$ | 0.4 ms |
| Low-pass $\tau_{LP}$ | 10 ms |

### Figure 4

In Figure 4 (Result 3), a Cox FPP is modeled using an AR(2) CIF, a GABA filter, and a low-pass filter where both filters are identical to those described in Results 1-2. In Panel A, the AR(2) CIF has noise variance *σ*^2^ = 0.315, and the mean rate of the process λ_0_ is 50 *spikes/sec*. Panel B keeps all parameters the same as Panel A, except the broadband component λ_0_ is increased to 400 *spikes/sec*. Panel C keeps all parameters the same as Panel A, except decreases the CIF (ensemble power) to *σ*^2^ = 0.1. All parameters are summarized in table 7.

**Table 7:** Figure 4 Model Parameters.

| Parameter | Value (Panel A) | Value (Panel B) | Value (Panel C) |
| --- | --- | --- | --- |
| GABA $\lambda_0$ | 50 *spikes/sec* | 400 *spikes/sec* | 50 *spikes/sec* |
| $\alpha_1$ | 1.9897557323136878 | unchanged | unchanged |
| $\alpha_2$ | -0.9901 | unchanged | unchanged |
| $\sigma^2$ | 0.315 | 0.315 | 0.1 |
| GABA $\tau_{decay}$ | 10 ms | unchanged | unchanged |
| GABA $\tau_{rise}$ | 0.4 ms | unchanged | unchanged |
| Low-pass $\tau_{LP}$ | 10 ms | unchanged | unchanged |

### Figure 5

In Figure 5 (Result 4), the CIFs of the subprocesses are shared up to a linear transform (given by Equation 27 with *w*_*A*_ = 1.5, *w*_*G*_ = −0.8, 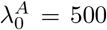, and 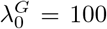), where the shared process (λ(*t*)) is a spectrally-define Gaussian process with a rhythmic peak at 1 Hz and peak power of 5000 *spikes/sec*. An additional low-pass filter is also used, as with Figure 2. The filters are given as *τ*_rise,GABA_ = 0.4 ms and *τ*_decay,GABA_ = 10 ms and *τ*_rise,AMPA_ = ms and τ_decay,AMPA_ = 4 ms. The low-pass filter is defined with *τ*_*LP*_ = 10 ms. The resulting power spectrum captures long timescale dynamics. All parameters are summarized in table 8.

**Table 8:**
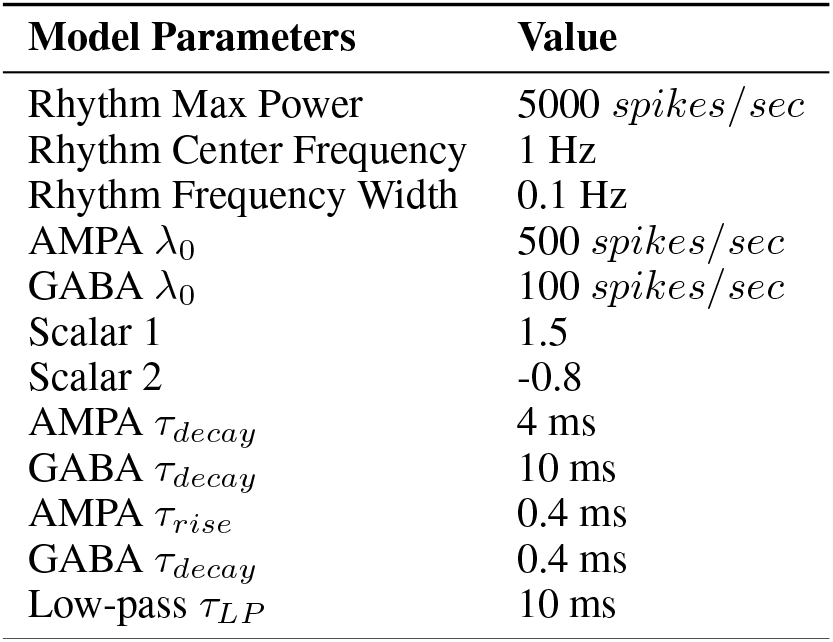
Figure 5 Model Parameters.

| Model Parameters | Value |
| --- | --- |
| Rhythm Max Power | 5000 *spikes/sec* |
| Rhythm Center Frequency | 1 Hz |
| Rhythm Frequency Width | 0.1 Hz |
| AMPA $\lambda_0$ | 500 *spikes/sec* |
| GABA $\lambda_0$ | 100 *spikes/sec* |
| Scalar 1 | 1.5 |
| Scalar 2 | -0.8 |
| AMPA $\tau_{decay}$ | 4 ms |
| GABA $\tau_{decay}$ | 10 ms |
| AMPA $\tau_{rise}$ | 0.4 ms |
| GABA $\tau_{decay}$ | 0.4 ms |
| Low-pass $\tau_{LP}$ | 10 ms |

Short timescale spectra are captured by generating a realization of the CIF for each subprocess. Conditioning on that realization λ(*t*), semi-stationarity can be assumed within small windows, so that the point process power spectra *S*_*X,A* |_ λ_*A*_(*t*) and *S*_*X,G*|_λ_*G*_(*t*) are approximately homogeneous, and equal to the rate at that time (λ_*A*_(*t*) and λ_*G*_(*t*), respectively). Conditioned on the rate, the two point processes are independent. After applying the postsynaptic filters and the low-pass filter, the cross-spectrum of the components is:

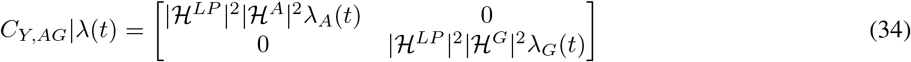

and the total power spectrum is:

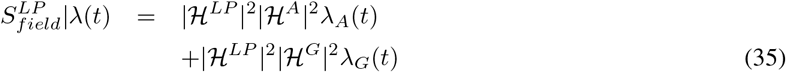

Panels B and C show the total spectrum and AMPA and GABA components (diagonals of Equation 34) for two chosen time points, and Panel E shows the total spectrum as a function of time (a spectrogram). Panel F shows the spectral slope and power at 80Hz as a function of time. Panel G shows the log ratio of the AMPA and GABA components from the spectrogram from E, i.e. log(*S*_*Y,A*_(*f*)/*S*_*Y,G*_(*f*)), where *S*_*Y,A*_(*f*) and *S*_*Y,G*_(*f*) are the diagonals of Equation 34.

### Implementation

The FFP framework, figure Jupyter notebooks, and all models were implemented in custom Python code and are available on the publicly available github package ‘filtered-point-process’ https://github.com/Stephen-Lab-BU/filtered-point-process)–an actively managed software package that will continue to be expanded in available features and tutorials.

## Acknowledgments

We’d like to thank the following individuals for valuable feedback: John Tauber, Becky Belisle, Proloy Das, Mingjian Alex He, Mark Kramer, and Uri Eden. We’d also like to thank the following funding opportunities: (Stephen) the Boston University Department for Mathematics and Statistics and the Boston University Center for Systems Neuroscience, (Oyama) the Boston University Undergraduate Research Opportunities Program, and (Bloniasz) NIH NINDS JSTPN T32NS131178 in early stage neuroscience, the NSF GRFP (2234657), and the Graduate Program for Neuroscience (GPN).

## Supplement 1: Methods for the data example in Figure 1

### Monkey ECoG data

In Figure 1, we analyze and model *Macaca fuscata* monkey ECoG data recorded under propofol anesthesia, where electrode coverage was over the left hemisphere (frontal, parietal, temporal, and occipital lobes). The data (M1) is documented and available in the public server Neurotycho.org [160] and is described experimentally in [75] and surgically in [160]. The monkey was recorded with a sampling rate of 1 kHz for 20 minutes while awake before being infused with 5.2 mg/kg of propofol until loss of behavioral consciousness (LOC). LOC was defined as the time when the monkey did not respond to manipulation of the hand or touching the nostril or philtrum with a cotton swab. Once LOC was established, data were recorded while the monkey was receiving propofol for 10 minutes before the infusion was stopped and the monkey recovered.

### Multitaper spectral analysis

A spectrotemporal estimate of 72 minutes of the recording session was generated using multitaper spectral estimation [161], with time resolution of 30 seconds and a frequency resolution of 2 Hz; the resulting spectrograms were averaged across electrodes (top Figure 1 Panel A). To highlight representative spectra as a function of brain state, an awake average spectrum is shown between 22 and 26 minutes (bottom left Figure 1 Panel A) and a behaviorally unconscious average spectrum is shown between 45 and 49 minutes (bottom right Figure 1 Panel A).

### FPP models of propofol anesthesia-induced unconciousness

The FPP framework was used to generate a forward model of the empirical spectrogram shown in Figure 1 Panel A. Three models were generated, ‘awake,’ ‘anesthetized,’ and ‘infusion’. As in the original study [75], we focus on the relatively stable dynamics in the ‘awake’ and ‘anesthetized’ states.

Awake and anesthetized states were modeled as a bivariate Cox process with AMPA and GABA subprocesses that share a CIF up to a scalar vector, as derived in Equation 28; the additional 1/f-like filter was also used, which distributes across Equation 28 similar to that shown in 25. The CIF is given as a spectrally defined Gaussian process, as in 29, with different rhythmic peaks in each state. The anesthetized model has two narrowband rhythms to capture the well-established slow wave and alpha rhythm. The awake model has a single ‘eyes closed’ alpha rhythm, but no slow wave.

The ‘infusion’ state was modeled as independent univariate homogeneous Poisson processes modeling AMPA and GABA with an additional 1/f-like filter.

The parameters for the ‘awake’, ‘infusion’, and ‘anesthetized’ models were chosen by hand-tuning (see Supplemental tables 1-3) so that the overall theoretical spectrum matched the empirical spectrum according to the mean absolute error, while constraining the parameters to create (1) rhythms in the slow and alpha range, and (2) AMPA and GABA waveforms within physiologically realistic ranges. Final model parameters are given in tables 1 (Awake), 2 (Anesthetized), and 3 (Infusion).

## Supplement 2: Model Parameter Tables

## Supplement 3: Asymptotic (Limiting) Slope Analysis

In this supplement, we perform an asymptotic analysis of the model structure shown in Figure 2 Panel I, focusing on the spectral slope as frequency *f* → *∞*. We start with the functional form of each filter and end with the limiting slope of the power spectrum for each model component.

### 1. Postsynaptic Potential

The Fourier Transform of the Postsynaptic Potential (PSP) filter is given by:

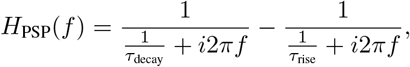

where *τ*_decay_ and *τ*_rise_ are time constants, and 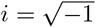.

#### Step 1.1: Simplify Each Term at High Frequencies

Define:

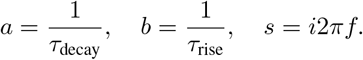

Then, the functional form becomes:

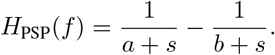

Compute the difference:

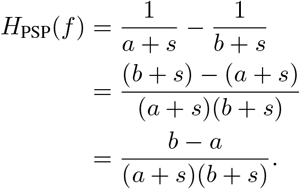

At high frequencies (*f* → ∞), |*s*| ≫ *a, b*, so:

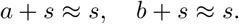

Thus, the denominator simplifies to:

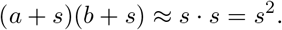

Therefore, the functional form simplifies to:

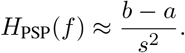

#### Step 1.2: Compute the Magnitude and Square the Expression

Compute the magnitude of *H*_PSP_(*f*):

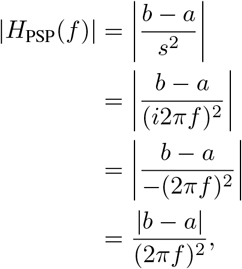

since *i*^2^ = −1 and the magnitude of a real number is its absolute value.

Then, compute the magnitude squared:

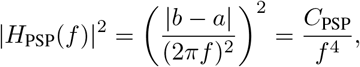

where:

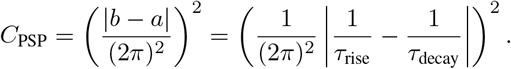

#### Step 1.3: Log-Log Slope

Taking the logarithm:

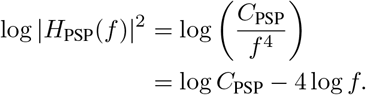

Here, we use the logarithm property 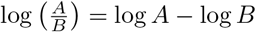 and log *f*^4^ = 4 log *f*.

**Impact on Slope:** The term log *C*_PSP_ is a constant with respect to *f*. **In a log-log plot, the slope is determined by the coefficient of** log *f*. Therefore, the constant term does not affect the slope.

Thus, the slope of |*H*_PSP_(*f*)|^2^ in the log-log domain is:

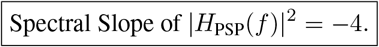

### 2. Low-Pass Filter (LP)

The low-pass filter functional form is given by:

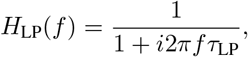

where *τ*_LP_ is the time constant of the low-pass filter.

#### Step 2.1: Simplify at High Frequencies

At high frequencies (*f* → ∞), the term *i*2*πfτ*_LP_ dominates the denominator:

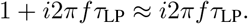

Therefore, the functional form simplifies to:

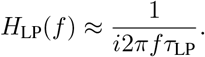

#### Step 2.2: Compute the Magnitude and Square the Expression

Compute the magnitude:

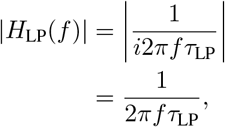

since |*i*| = 1.

Then, compute the magnitude squared:

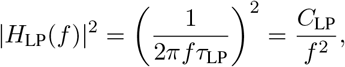

where:

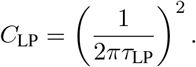

#### Step 2.3: Log-Log Slope

Taking the logarithm:

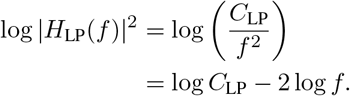

Here, we use 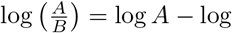 B and log *f* ^2^ = 2 log *f*.

**Impact on Slope:** The constant term log *C*_LP_ does not affect the slope. The slope is determined by the −2 log *f* term. Thus, the slope of |*H*_LP_(*f*)|^2^ in the log-log domain is:

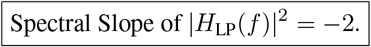

### 3. Point Process Spectrum

The Cox point process spectrum, which includes both rhythmic (e.g., Gaussian) and broadband (constant rate) components, is given by:

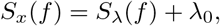

where *S*_*λ*_(*f*) represents the conditional intensity function (CIF), and λ_0_ is the rate of the process, representing a flat spectrum.

As an example of a CIF for the process, we use a spectrally defined Gaussian process where the shape of the power spectrum is defined as a Gaussian bump given by:

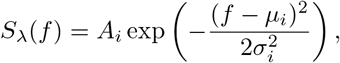

where *A*_*i*_, *µ*_*i*_, and *σ*_*i*_ are constants.

#### Step 3.1: Asymptotic Behavior at High Frequencies

As *f* → ∞, the Gaussian component *S*_*λ*_(*f*) decays exponentially to zero:

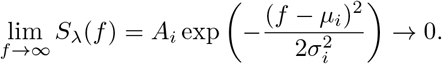

Therefore, at high frequencies, the Gaussian term *S*_*λ*_(*f*) contributes negligibly to the spectrum relative to the non-zero constant broadband component. However, the slope of *S*_*λ*_(*f*) will not be strictly zero in log-log.

#### Step 3.2: Log-Log Slope

*S*_*λ*_(*f*) decays (exponentially) faster than any linear rate in log-log space. As such, its limiting slope is:

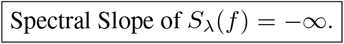

Even though the limiting slope is non-zero because of the exponential decay, the Gaussian component will not contribute to the overall model limiting slope because, relative to the broadband component, *S*_*λ*_(*f*) is not contributing much to the spectrum at high frequencies. Because the remaining term is a constant, the slope from λ_0_ is 0. Thus, the spectral slope of the point process spectrum *S*_*x*_(*f*) at high frequencies is:

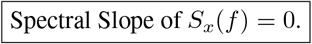

**Note:** It’s important to note that while *S*_*λ*_(*f*) decays to zero exponentially fast, λ_0_ remains constant. There are scenarios where the spectral slope of *S*_*x*_(*f*) might not be zero, which would occur when the rhythmic component *S*_*λ*_(*f*) would dominate over the broadband component λ_0_ at high frequencies (i.e., the spectrum decays at a rate slower than linear in log-log space).

### 4. Full Combined Model

The full combined model is:

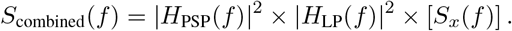

#### Step 4.1: Asymptotic Behavior at High Frequencies

Using the asymptotic forms from previous sections:

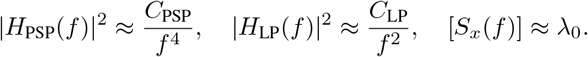

Therefore, at high frequencies:

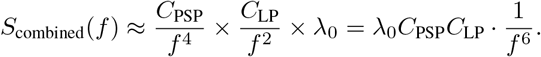

#### Step 4.2: Log-Log Slope

Taking the logarithm:

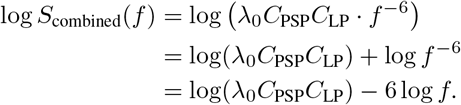

Here, we use the logarithm properties log(*AB*) = log *A* + log *B* and log *f* ^−6^ = −6 log *f*.

**Impact on Slope:** The constant term log(λ_0_*C*_PSP_*C*_LP_) does not depend on f and thus does not affect the slope. **The slope is determined solely by the** −6 log *f* **term**.

Thus, the spectral slope of the full combined model at high frequencies is:

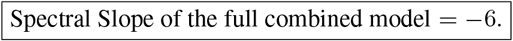

### Summary

This asymptotic analysis demonstrates that at high frequencies (*f* → ∞), the spectral slope of the full combined model is determined by the sum of the slopes of the individual filter components and the point process spectrum. Specifically:

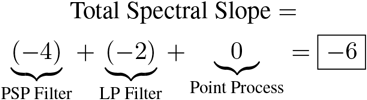

This style of limiting slope analysis can be applied to any set of model components that have an analytical solution (e.g., event/additional filters, point processes).

## Supplement 4: Full Gabor Fourier transform with scaling constant

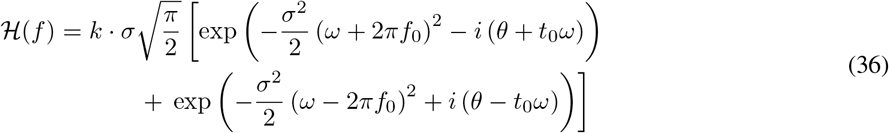

where

- *ω* = 2*πf* – Angular frequency,
- *k* – Amplitude scaling factor,
- *f*_0_ – Center frequency,
- *σ* – Standard deviation of the Gaussian envelope,
- *θ* – Phase offset,
- *t*_0_ – Time shift.

